# Integrated single-cell analysis defines the epigenetic basis of castration-resistant prostate luminal cells

**DOI:** 10.1101/2023.03.03.530998

**Authors:** Jason S Kirk, Jie Wang, Amanda Tracz, Mark Long, Spencer Rosario, Yibing Ji, Rahul Kumar, Xiaozhuo Liu, Prashant K Singh, Igor Puzanov, Gurkamal Chatta, Qing Cheng, Jiaoti Huang, Jeffrey L Wrana, Jonathan Lovell, Han Yu, Song Liu, Michael M Shen, Tao Liu, Dean G Tang

## Abstract

Understanding prostate response to castration and androgen receptor signaling inhibitors (ARSI) is critical to improving long-term prostate cancer (PCa) patient survival. Here we use a multi-omics approach on 229,794 single cells to create a mouse single-cell reference atlas better suited to interpreting mouse prostate biology and castration response. Our reference atlas refines single-cell annotations and provides chromatin context, which, when coupled with mouse lineage tracing demonstrates that the castration-resistant luminal cells are distinct from the pre-existent urethra-proximal stem/progenitor cells. Molecular pathway analysis and therapeutic studies further implicate JUN/FOS, WNT/*β*-Catenin, FOXQ1, NF*κ*B, and JAK/STAT pathways as the major drivers of castration-resistant luminal populations with high relevance to human PCa. Importantly, we demonstrate the utility of our datasets, which can be explored through an interactive portal (https://visportal.roswellpark.org/data/tang/), to aid in developing novel combination treatments with ARSI for advanced PCa patients.

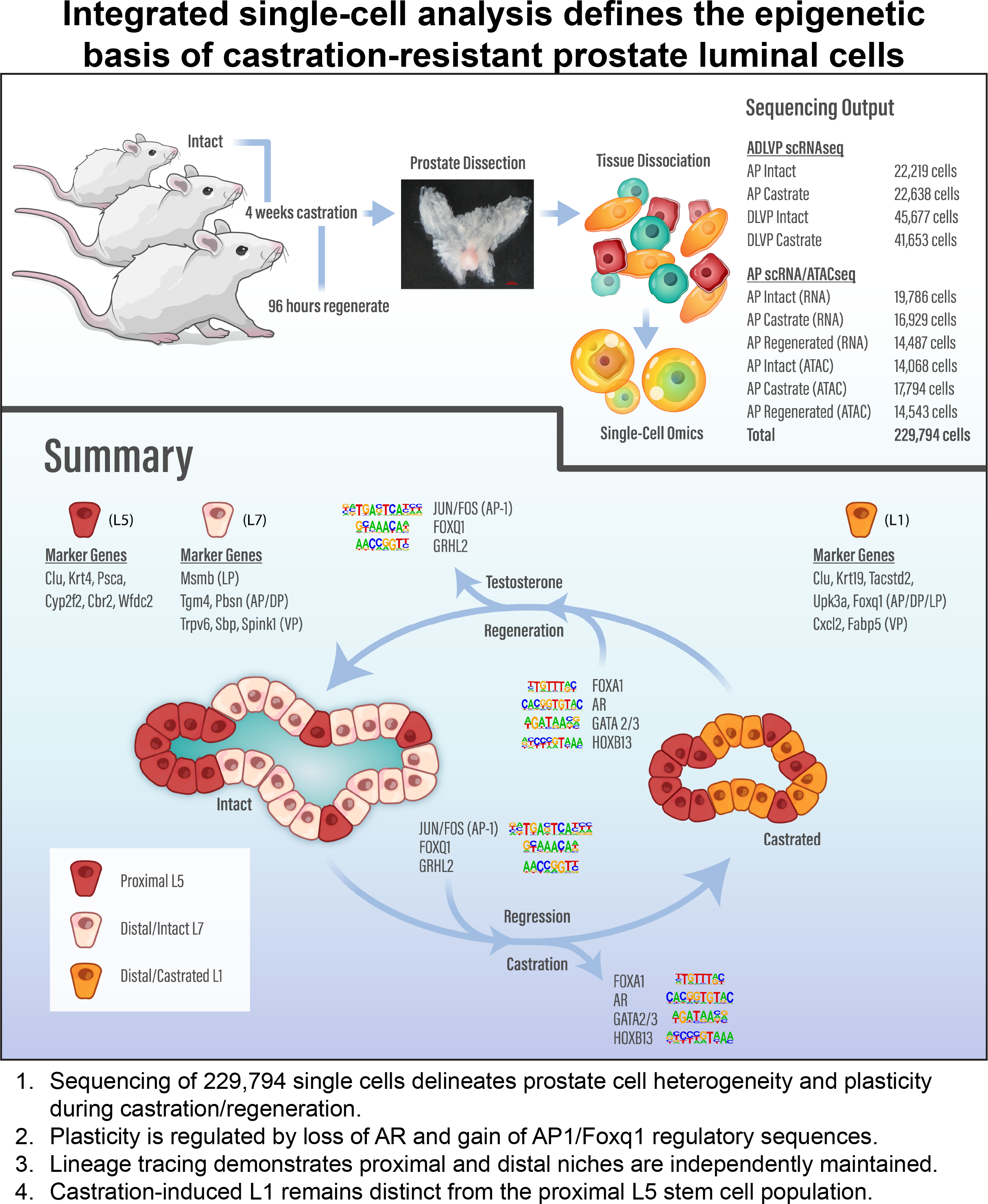

## INTRODUCTION

The prostate is an exocrine gland that secretes proteins into the ejaculatory duct to support reproductive success. Traditionally, the prostate was simply thought to be composed of luminal, basal, and scarce neuroendocrine epithelium surrounded by supportive stroma^1^. However, recent single-cell RNA sequencing (scRNAseq) studies have revealed a more complex gland comprised of numerous functional and supporting cell subtypes^2–9^. All cell populations contribute to the homeostatic functions of the prostate, but oncogenic transformation in epithelium and disruption of glandular homeostasis result in prostate cancer (PCa).

PCa is the most diagnosed and the second leading cause of cancer-related mortality in the U.S^10^. Although low-grade disease has favorable treatment outcomes, men diagnosed with locally advanced and metastatic disease have a poor prognosis. Androgen deprivation therapy (ADT) combined with radiotherapy is standard of care for advanced PCa patients, and addition of androgen receptor (AR) signaling inhibitors (ARSIs) such as enzalutamide may offer added benefits^11^. Given the prostate’s reliance on androgen signaling for growth and differentiation, ARSIs can be highly effective. Unfortunately, most advanced PCa patients eventually become refractory to ARSIs and develop lethal castration-resistant PCa (CRPC) or metastatic CRPC (mCRPC)^12^. It is critical that we understand how prostate cells respond to castration and develop resistance so that we can design better therapeutic approaches for PCa. Some mechanisms of resistance are well established^13^, but mechanisms related to epigenetic plasticity are just beginning to be appreciated^14^.

The rodent has been classically used to study prostate response to castration. Androgen removal via castration results in reduction of prostate weight due to apoptosis of a high percentage (as much as 90%) of glandular luminal cells^15–17^. Importantly, addition of exogenous androgens results in complete regrowth and differentiation of the atrophied prostate^18–20^. These and other seminal discoveries led to the postulation that pre-existent, castration-resistant, stem cells were the source of prostate regeneration^21^. It is known that early during prostate development (and perinatally), primitive stem cells are concentrated in the prostatic buds proximal to the urethra, which undergo branching morphogenesis to generate the distal tubules and ducts^22, 23^. In the adult mouse prostate, dormant stem cells are also most abundantly observed in the proximal prostate^24–28^ although both basal and luminal cell compartments are largely self-sustained by their own progenitor cells scattered throughout the distal prostate^29, 30^. It was thought that proximally located luminal stem/progenitors are what give rise to regenerated glands after successive rounds of castration/regeneration. However, a recent scRNAseq study suggested that both distal and proximally located luminal cell populations have equal potential to regenerate prostates and that during castration differentiated luminal cells undergo reprogramming to become proximal-like progenitors and then castration-resistant^8^. Whether these two cell populations completely converge and where the reprogrammed luminal cells originate from remain unclear. Do all differentiated luminal cells have the potential for reprogramming, or is it a specific subset of luminal cells? Answers to these questions and further understanding the biology underlying prostate response and resistance to castration could have important implications for determining clinical response and resistance to ARSIs.

Using both scRNA and scATAC sequencing of 229,794 mouse prostate cells, we have created a unique reference atlas capable of more completely understanding the biological processes contributing to the castration/regeneration response. Our robust dataset demonstrates that castration- reprogrammed luminal cells remain distinct from the pre-existent proximal progenitors. Lineage tracing using proximally and distally restricted models further demonstrates that proximal luminal progenitors do not give rise to distal glands, and that distal luminal cells can survive and regenerate the gland after castration. Notably, integrating scRNAseq with single-cell chromatin landscapes link JUN/FOS and FOX family members as key regulators of castration response. Lastly, pathway analysis of castration response reveals numerous therapeutic targets and clear relationships with CRPC development in patients. These findings serve as a resource (https://visportal.roswellpark.org/data/tang/) to future investigators and may contribute to new therapeutic development for men who have failed ADT and ARSIs.

## RESULTS

### scRNAseq identifies castration-resistant populations in all 3 major cellular compartments

To better understand castration response, we harvested intact and castrated wild-type C57/BL6J mouse prostates (Figure S1A-C) for scRNAseq analysis (Figures 1, S1-S3). <u>D</u>orsal, lateral, and <u>v</u>entral <u>p</u>rostate (DLVP) lobes and <u>a</u>nterior <u>p</u>rostates (AP) were analyzed as separate cohorts. Upon processing and quality control (see Methods), we identified 132,187 high-quality cells across 4 sample cohorts (Figure S1D; Table S1), which were classified into 23 distinct cell clusters (Figures 1A-C, S2B). Clusters included 8 luminal (L1-L8; *Krt8/18^+^, Cd24a^+^, Epcam^+^, Krt5/14^-^*), 3 basal (B1-B3; *Krt5/14^+^, Tp63^+^, Lgals7^+^*), 4 mesenchymal (Me1-4; *Vim^+^, Col1a1^+^, Igf1^+^*), and multiple immune and supporting stromal cell populations (Figure S2). Too few neuroendocrine (NE) cells were identified to form an independent cluster (not shown).

**Figure 1.**
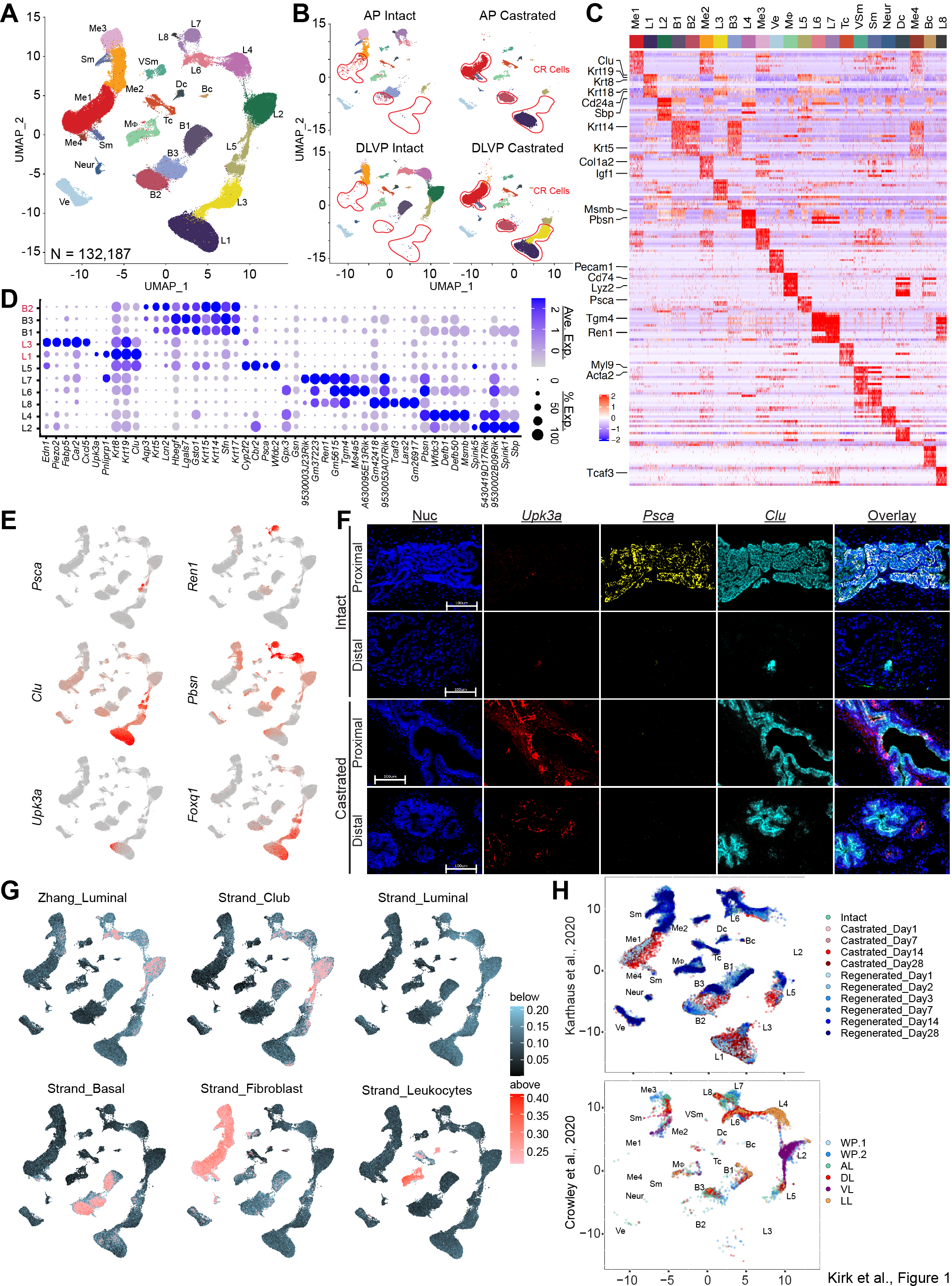
Castration induces cell population changes in the 3 major cell lineages of prostate. (**A-B**) UMAP of scRNAseq in 132,187 cells of the mouse prostate under intact and castrated conditions. Shown in B are scRNAseq UMAPs of individual sample libraries from intact and castrated mice. Red boundaries indicate castration-resistant populations. L, luminal; B, basal; Me, mesenchymal; Sm, smooth muscle; VSm, vascular Sm; M*Φ*, macrophage; Tc, T cells; Bc, B cells; Dc, dendritic cells; Ve, vascular endothelium; Neur, neural. (C) Heatmap of cell-specific genes identified from scRNAseq with annotation of lineage specific markers. (D) Bubble plot of unique marker genes from epithelial cell subtypes. (E) Representative marker gene expression overlayed on UMAP projection demonstrates cluster specificity. (F) Representative images from *in situ* RNA hybridization of luminal-specific markers in intact and castrated prostates examined at proximal and distal regions. Scale bar, 100 μm. (G) Human cell type-specific gene signatures mapped to single cells using AUCell from SCENIC. Cells with specific correlation enrichment above threshold are shaded red according to enrichment score. (H) Reference based remapping of Karthaus *et al.^8^* and Crowley *et al.^3^* datasets to the ADLVP single cell reference atlas, color-coded by library sample subtype.

Under intact conditions, AP was composed of 4 unique cell populations, Me3, B3, L7 and L8. Me3 uniquely expressed *Mmp2, Mfge8*, and *Wif1* (Figure S1I) and B3 *Aqp3*, *Areg*, and *Dusp7* (Figure 1D; Table S2). L7 and L8 were characterized by expression of *Ren1* and *Tcaf3,* respectively (Figures 1D-E). The DLVP showed 4 unique cell populations, B1, L2, L4 and L6, with L2, L4, and L6 associated with VP, LP, and DP, respectively (Figure S2A) and characterized by high expression of *Sbp*, *Msmb, and A630095E13Rik* transcripts, respectively (Figures 1D, S2E). Consistent with earlier studies^31^, the VP lacked *Pbsn* (Figures 1D-E), which was confirmed by whole-mount *in situ* hybridization (Figure S1F). The L5 cell population, characterized by *Psca, Wfdc2, Clu,* and *Cyp2f2* expression, was shared between AP and DLVP (Figures 1D-F, S2F,H). RNAscope using *Pbsn* and *Clu* probes demonstrated that L5 was proximally located relative to L7 in the AP (Figure S1E).

After 4 weeks of castration, there was apparent loss of differentiated luminal cells (L2, L4, L6, L7, and L8) with simultaneous emergence of two *de novo* luminal populations, L1 and L3 (Figures 1A- D). L1, the major castration-resistant luminal population, was defined by high levels of *Clu, Upk3a, Krt19, Foxq1* and *Krt18* expression (Figures 1D, S2, S3F). Unlike proximal L5, L1 *does not* express *Psca, Cyp2f2, Cbr2,* or *Wfdc2*, marking L1 as a unique population arising due to castration. L3 was defined by expression of *Edn1, Piezo2, Fabp5,* and *Cxcl5* but also expressed high levels of *Clu, Krt19* and *Krt18* (Figures 1D-E, S2D,F). Expression of *Upk3a* and *Clu* were validated in AP by RNAscope (Figure 1F), and *Upk3a* was only expressed after castration (Figures 1E-F, S2G). Conversely, *Clu* was detected proximally in L5 cells under intact conditions and further induced distally post castration (Figures 1E-F, S1E-F). As in previous studies^3, 4, 6, 8^, scattered L5 cells, as evidenced by *Psca* and *Clu* expression, were found in the distal prostate (Figures 1E-F, S1E-F).

Castration also induced a dramatic shift in both mesenchymal (Me) and basal (B) cell populations. Me3 and Me2 disappeared with castration while Me1 became predominant (Figures 1A-B). Castration- surviving Me1 was featured by loss of differentiation markers *Col3a1, Col1a2* and *Gsn* while gaining *Apod* and *Apoe* (Figure S1I; Table S2). Me4 was also identified with castration but was suspected to be a doublet and not examined further. Within the basal compartment, B1 and B3 were lost with castration, giving rise to B2, which showed upregulation of *Aqp3, Lcn2* and *Krt5* and downregulation of *Areg* and *Apoe* (Figures 1D, S2J; Table S2). All other cell populations were minimally perturbed by castration (Figure S1I), demonstrating the specificity of androgen signaling in regulating the 3 major prostate cell lineages.

Using AUCell from SCENIC^32^, we correlated the individual cells of our ADLVP dataset (Figures 1A-B) to relevant human gene signatures^5, 33^, and observed strong concordance between the corresponding populations (Figure 1G; Table S7, see below). Single-cell gene signatures from Henry *et al.*^5^ demonstrated that human basal, fibroblast, leukocyte, and club cell signatures all highly correlated with corresponding murine populations (Figure 1G). Interestingly, luminal signatures in Zhang *et al.*^33^ correlated well to ADLVP but luminal cells from Henry *et al.*^5^ did not, likely due to differences in luminal-specific genes between mouse and human.

### ADLVP single-cell data serves as a reference atlas to better define prostate single-cell datasets

To our knowledge, no other study to date has interrogated as many prostate cells under intact and castrated conditions as this study (Figure S2C). We hypothesized that our ADLVP dataset, together with scATACseq data (below) could better serve as a reference atlas to map cell states^34^. To test this hypothesis, we utilized our ADLVP dataset as a reference atlas to remap two previous studies^3, 8^ (Figures 1H, S3A-D). Indeed, remapping of Karthaus *et al.*^8^ refined all cell populations across castration timepoints (Figures 1H, S3A,C,E) and demonstrated the loss of differentiated populations with concomitant emergence of castration-resistant populations. Karthaus *et al.*^8^ identified luminal_1, luminal_2Psca^+^, and luminal_3Foxi1^+^ populations, but reference mapping identified 5 distinct luminal populations (L1, L5, L6, L7, and L8). L5 reflected luminal_2Psca^+^ cells whereas Luminal_1 was refined into L6, L7, and L8 populations and luminal_3Foxi1^+^ most closely resembled L5 (Figure S3E). Reference mapping also showed that L1 was present but undefined as a unique population by Karthaus *et al*.^8^, likely due to relatively low cell numbers sequenced in that study. Notably, the reference remapping revealed that L1 and L5 remained distinct cell populations (Figures 1H, S3A,C,E). Lastly, cell score predictions showed how well other cell populations from Karthaus *et al.* matched annotated clusters from our ADLVP (Figure S3E). Crowley *et al.*^3^ focused on both the whole prostate and lobe-specific populations under intact conditions. We found that the lobe-specific luminal populations in Crowley *et al.*^3^ mapped to the exact coordinates of our reference map for corresponding luminal populations (Figures 1H and S3B, D). Again, previously unidentified populations were identified from the Crowley *et al.*^3^ dataset, namely, L8, B1 and B3. Overall, these results demonstrate the superior capability of our ADLVP dataset to serve as a reference atlas to better define future and refine past single-cell studies (Figure S2C).

### Integrated scRNA and scATAC sequencing further defines castration-specific cell subtypes

Castration induced several *de novo* castration-resistant cell populations, including L1, L3, B2, and Me1 (Figure 1). To elucidate the regulatory programs driving the generation of castration-resistant populations, we performed an integrated sequencing study combining transcriptomic and chromatin profiling at the single-cell level (Figures 2, S4, S5; Table S1). For simplicity, only AP was chosen for these studies and prostates were harvested from intact, castrated, and regenerated mice (Figure S4A). scRNAseq and single-cell Assay for Transposase-Accessible Chromatin sequencing (scATACseq) were processed in parallel from the same samples. After sequencing and processing, 51,202 high-quality cells from scRNAseq and 46,405 high-quality cells from scATACseq were identified (Figures 2A- B; Tables S3, S4). scRNAseq cell clusters were generated as above and designated with the same cluster annotations as in Figure 1. Interestingly, 2 additional luminal cell clusters, L9 and L10 were observed (Figure 2A). L9 was identified after regeneration whereas L10 was not previously delineated but seemingly present as part of the L7 cluster in the ADLVP study (Figure 1A). L10 and L7 overlapped significantly but L7 was distinguished by higher expression of *Sox9, Cxadr,* and *Itga6* (Figure S4E). L9 was defined by expression of *Tgm4* and *Pbsn* but not *Ren1* (Figure S4D). In addition to novel luminal populations, seminal vesicle luminal cell (SvL) contamination distinguished by expression of *Svs2*, *Svs4*, *Svs5*, and *Pate4*, as well as an additional mesenchymal population Me5 were detected (Figures 2A, S4D). Like L9, Me5 was specific to regeneration. Me5 was very similar to Me3 but found to have higher absolute levels of Me3 target genes. For example, Me3 expressed *Col1a2* at 0.66 avg_logFC while Me5 at 1.66 avg_logFC (Table S3). These differences could be due to an above average response to replenished testosterone after castration. Importantly, two independent experiments (i.e., ADLVP and AP2; Table S1) have now observed the same castration-induced cell population shifts (Figures 1B, 2C). All other ‘accessory’ cell types in the prostate showed only minor changes with castration. Significantly, independent scRNAseq experiments also revealed the L1 and L5 to be distinct cell populations (Figures 2A, S4F,G), and remapping of the Karthaus *et al.*^8^ dataset with only the AP dataset as a reference atlas again demonstrated these dynamic changes and distinctions (Figure S4H).

**Figure 2.**
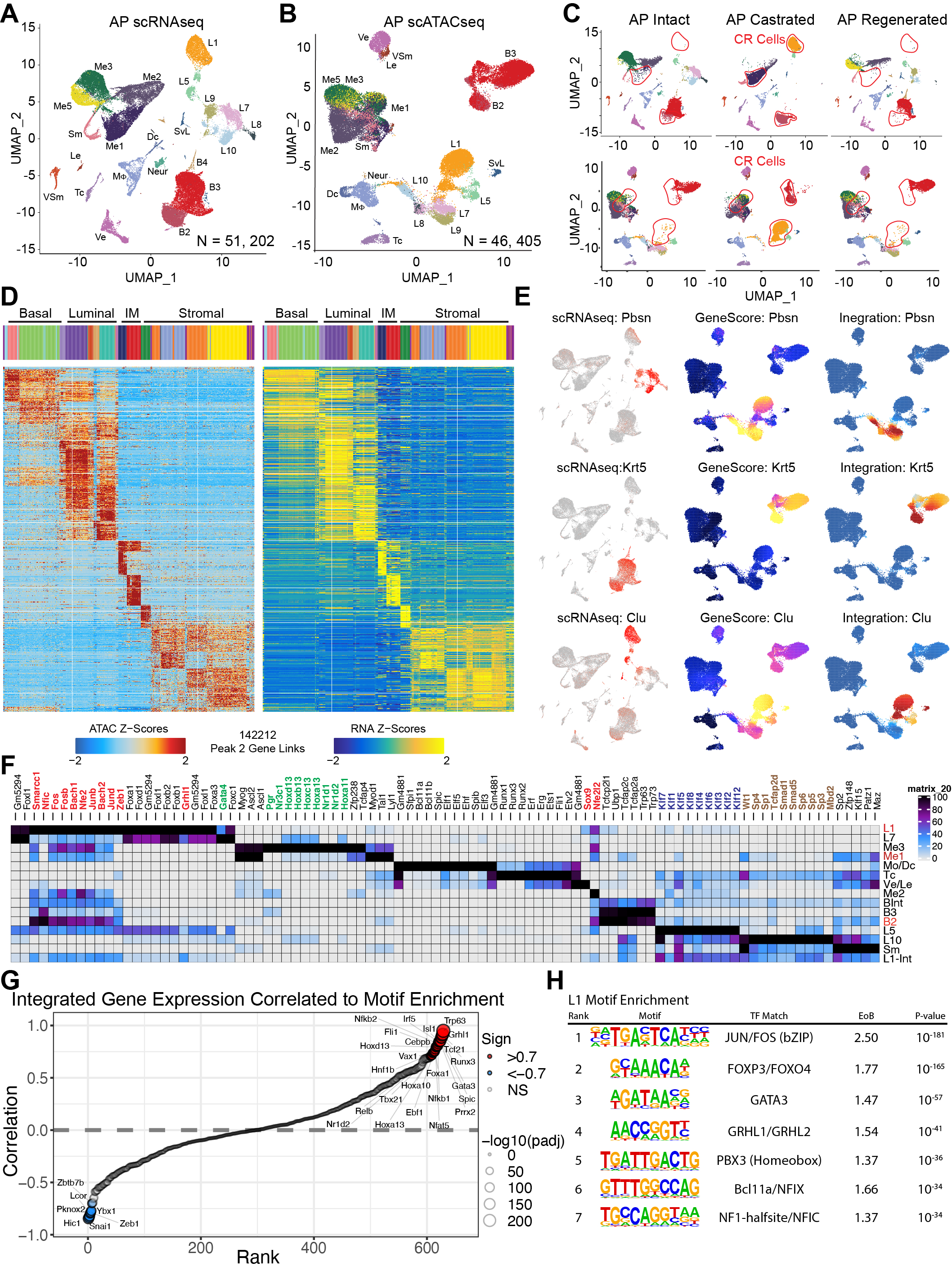
Integrated scRNAseq and scATACseq analysis identifies key mediators of prostate castration response. **(A-C)** UMAP of the AP scRNAseq (A) and scATACseq (B) data with cell clusters annotated as above. Shown in (C) are library-specific UMAPs identifying castration-resistant (CR) cell populations (red boundaries). Le, lymphatic endothelium; SvL, seminal vesicle epithelium (D) Peak to gene (P2G) linkage showing the correlations between ATACseq peak accessibility and RNAseq expression profiles for different cell populations. (E) Lineage restricted examples of scRNAseq and scATACseq data integration, represented by *Pbsn, Krt5,* and *Clu genes*. (F) Heatmap of the known motifs enriched in open chromatin of intact and castrated single-cell clusters. (G) Rank ordered plot of known motifs most positively and negatively correlated to gene expression from integrated single-cell datasets. (H) *De novo* motif sequence enrichment in L1 specific chromatin peaks ranked by significance. EoB, Enrichment over background.

Next, we asked whether the chromatin landscape (Figure S5) could shed molecular insights into the biology driving castration response. Using ArchR^35^, the scATACseq identified 16 high-quality cell clusters distinguished by chromatin accessibility (Figures S5A-C). Since ATACseq is highly enriched in transcription start sites (TSS), these regions, along with chromatin accessibility across gene bodies, were used to predict gene activation, i.e., GeneScores (Figure S5D; Table S4). For integration purposes, the GeneScore matrix from scATACseq was correlated to gene expression matrix from the scRNAseq dataset. Integration with ‘labeltransfer’ provided cell cluster annotation of the scATACseq populations matching those of the scRNAseq dataset (Figure 2B). Post integration, 142,212 chromatin peaks were matched to corresponding genes from the scRNAseq dataset using Peak 2 gene integration (Figure 2D). To confirm proper integration, a pool of population-specific markers was used and found to have high levels of concordance between gene expression and predicted GeneScores (Figure 2E). Chromatin accessibility of lineage-specific genes such as *Pbsn* was found to be limited to differentiated luminal cells (Figure S5H). There was residual accessibility in castration-resistant L1, yet one accessibility peak near the gene promoter was reduced, suggesting decreased transcriptional output (Figure S5H, region A). Conversely, *Clu* chromatin accessibility was restricted to proximal L5 and castration-resistant L1 populations (Figure S5H, region B). As with *Pbsn*, there was residual *Clu* gene accessibility in L7/L9 population that likely came from L9 cells, which were found in 96-hr regenerated prostates and probably had not completely closed chromatin around this castration-induced gene. Collectively, the scATACseq analysis reaffirmed our scRNAseq findings, and the integration was highly successful allowing confident identification of scATACseq cluster populations.

### JUN/FOS and FOX family motifs are the major regulatory factors in castration-resistant L1

Analysis of known transcription factor (TF) motif enrichment identified distinct transcriptional programs under intact and castrated conditions (Figure 2F; Table S5). Surprisingly, although AR (*Nr3c4*) was deregulated by castration, it was not identified in the top deregulated motifs. In contrast, fellow nuclear receptor family members *Nr3c1* (encoding glucocorticoid receptor or GR) and *Nr3c3* (*Pgr*) that share a conserved DNA binding motif were observed (Figure 2F). Enrichment for *Ar*, *Nr3c1*, and *Pgr* motifs was most apparent in differentiated Me3/Me2 populations followed by intact L7, intact B3, and proximal L5 cells, but negligible in L1 and other cell populations. As expected, enrichment for all NR3C family members was diminished with castration across responsive populations. In addition to *Nr3c* motifs, *Fox/Gata/Hox* family motifs were enriched in luminal cells whereas the *Trp63* motif was enriched in basal cells (Figure 2F). Strikingly, the proximal L5 was highly enriched in *Klf, Tead3/4, Zeb1, and Fox* family motifs although *Runx2* motifs were also observed at low levels. In contrast, *Gata* motifs were found to be highly enriched in differentiated luminal cells (L7/L8/L9/L10) but not in proximal L5 (Figure 2F), highlighting key differences between the proximal and distal regions. In addition to AR regulation, mesenchymal populations (Me1/2/3) were enriched in *Ascl1/2*, *MyoG*, and *Tal1* motifs. Macrophages and dendritic cells were enriched for *Bcl11a/b*, *Spib/c* and *Elf* family motifs whereas T cells in *Runx* and *Erg/Ets* motifs (Figure 2F). Further dissecting the list of regulatory motifs, we identified the motifs most significantly correlated to integrated gene expression (Figure 2G). Unsurprisingly, lineage-restricted motifs such as *Trp63, Tcf21, Gata3* and *Foxa1* were significantly and positively correlated with gene expression. Motifs negatively correlated with gene expression were not likely to be functionally relevant since they would be expressed/regulated in different cell populations.

In general, we observed a clear loss of differentiation-regulatory motifs with castration (e.g., *Ar, Gata, Hox*) and a simultaneous enrichment for stem cell-regulating motifs (e.g., *Zeb1, Klf* and *Sox9*) (Figure 2F; Table S5). Strikingly, L1 cells were most notably enriched in *Fox, Grhl* and *Jun/Fos* motifs (Figure 2F). As searching for known TF motifs causes a lot of overlapping family members to be found (even when not expressed), we performed *de novo* motif analysis using HOMER^36^, which could distil regulatory elements down to exact motif sequences, and identified the top regulatory elements for each cell cluster (Figures 2H, S5E-G). Significantly, *de novo* analysis of L1 similarly identified *Jun/Fos*, *Fox* and *Grhl* motifs as the top regulatory factors in addition to several others including *Gata* motifs (Figure 2H). Interestingly, differentiated L7 was also enriched in *Fox*, *Gata* and, to a lower extent, *Ets*, *Jun/Fos*, *Hox*, and *Ar* family motifs (Figure S5E). Proximal L5 was enriched for motifs representing *Fox*, *Jun/Fos*, *Klf* and *Elf3* family members (Figure S5F). Lastly, the least well-defined transitory L9 population was found to be enriched for *Maz, Osr2, Nfat, Jun/Fos* and *Arid3b* motifs (Figure S5G).

Taken together, the chromatin landscape and motif analysis identified *Jun/Fos* (AP-1), *Fox, and Grhl* family motifs as the main regulatory factors driving castration resistance and demonstrated that L1 and L5, although both castration-resistant, represent 2 epigenetically distinct cell populations.

### Lineage tracing confirms that L1 does not originate from the pre-existent L5 population

Karthaus *et al*.^8^ suggested that castration caused differentiated luminal cells to become L5-like cells which then gave rise to castration-resistant (L1) cells. However, our 2 batches of scRNAseq in >180,000 cells (Figure S2C; Table S1), whole-mount RNAScope (Figures 1F, S1E-F) and scATACseq studies indicated that L1 and L5 populations were distinct, and L5 did not expand upon castration. We employed several mouse lineage-tracing models (Figures 3-4, S6) to further elucidate the L1 cell origin and determine whether L5 expands in response to castration and gives rise to L1 during castration/regeneration. All lineage tracing models were analyzed along the same timeline used for scRNAseq/scATACseq studies (Figures 3A, 4A). We first utilized a *Clu* driven Cre-ER(T2);tdTomato (*Clu;*Tomato) model^37^, since under intact conditions *Clu* was restricted to proximal prostate (Figures 1D, S4D) whereas under castrated conditions both proximal L5 and distal L1 cells expressed *Clu* (Figures 1E-F, S1E-F). Consistently, intact *Clu;*Tomato mice showed prominent Tomato expression in proximal AP with only sparse labeling of distal luminal cells (Figure 3B). Remarkably, in castrated and regenerated prostates, Tomato was still predominantly expressed in proximal niche with no expansion in either proximal or distal prostates (Figures 3C, S6A). Interestingly, when Tomato was activated *following* castration and *after* the *de novo Clu^+^* L1 population emerged, all cells in proximal and distal regions were labeled with Tomato (Figures 3D, S6B). Quantification of Tomato^+^ KRT8 and KRT5 expressing cells showed that Tomato^+^ cells were mainly located in KRT8^+^ luminal layer of the intact proximal prostate, and not expanded with castration (Figures 3E-F).

**Figure 3.**
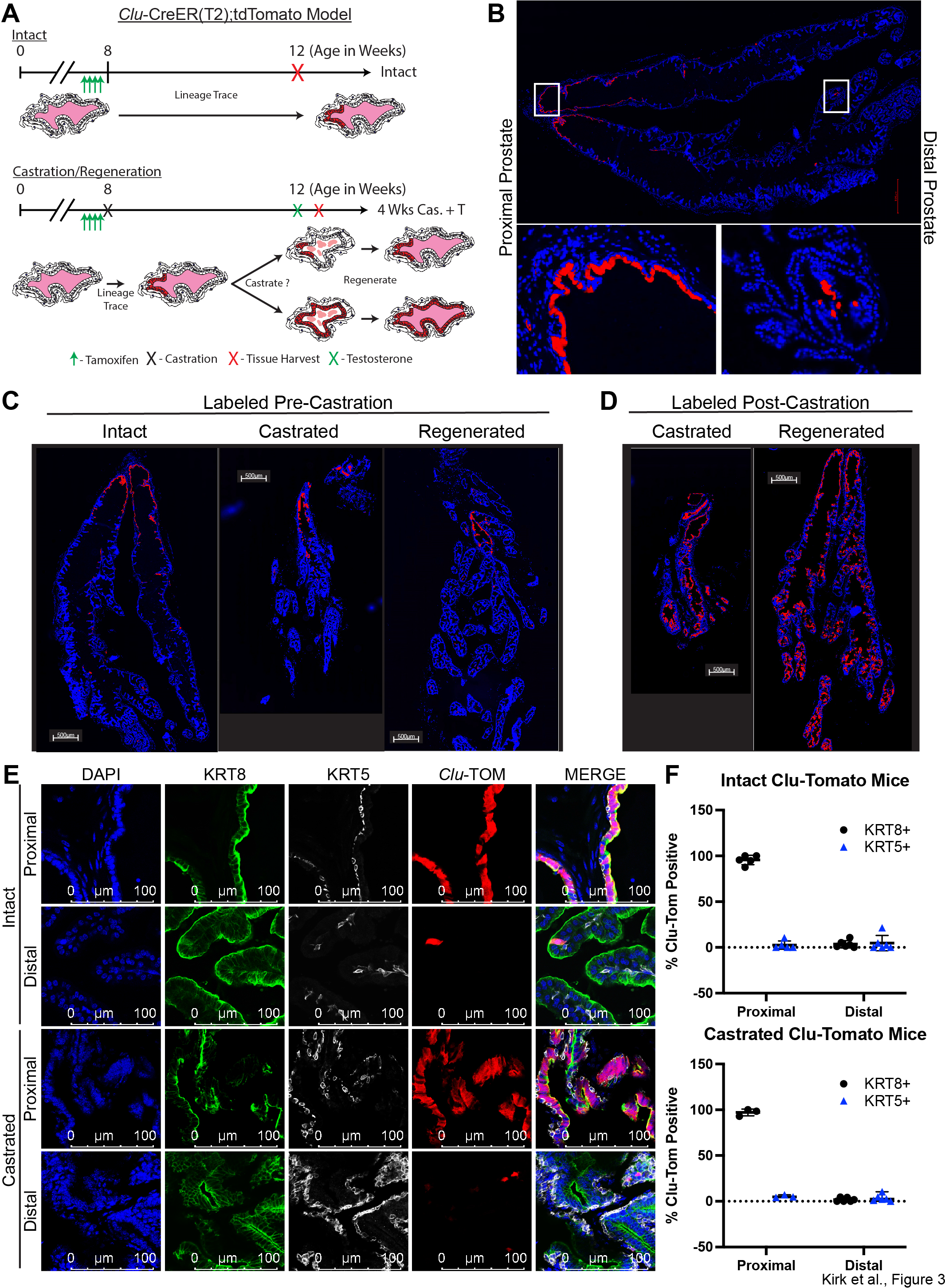
*Clu-*CreER(T2) lineage tracing confirms L1 does not originate from the L5 population. **(A)** Schematic demonstrating expected outcomes of lineage-tracing studies when Tomato tracer is activated by Tamoxifen in intact and castration/regeneration settings. **(B)** In intact prostate, lineage tracing ubiquitously labeled proximal luminal cells (red) while sparsely labeled distal luminal cells. White boxed regions are enlarged (below). (**C-D**) Representative whole-mount images of pre-castration (C) or post-castration (D) labeled *Clu*;Tomato APs from intact, castrated, and regenerated prostates. (E) Representative images of pre-castration labeled *Clu*;Tomato AP double stained for KRT8 (AF-488) or KRT5 (AF-647) in proximal and distal regions of intact and castrated prostates. (F) Quantification of proximally and distally labeled Tomato^+^ luminal and basal cells from intact and castrated prostates. Scatter plots represent the mean ± SD of 3 independent regions of interest (ROI) in proximal and distal AP from 2 independently labeled mice.

**Figure 4.**
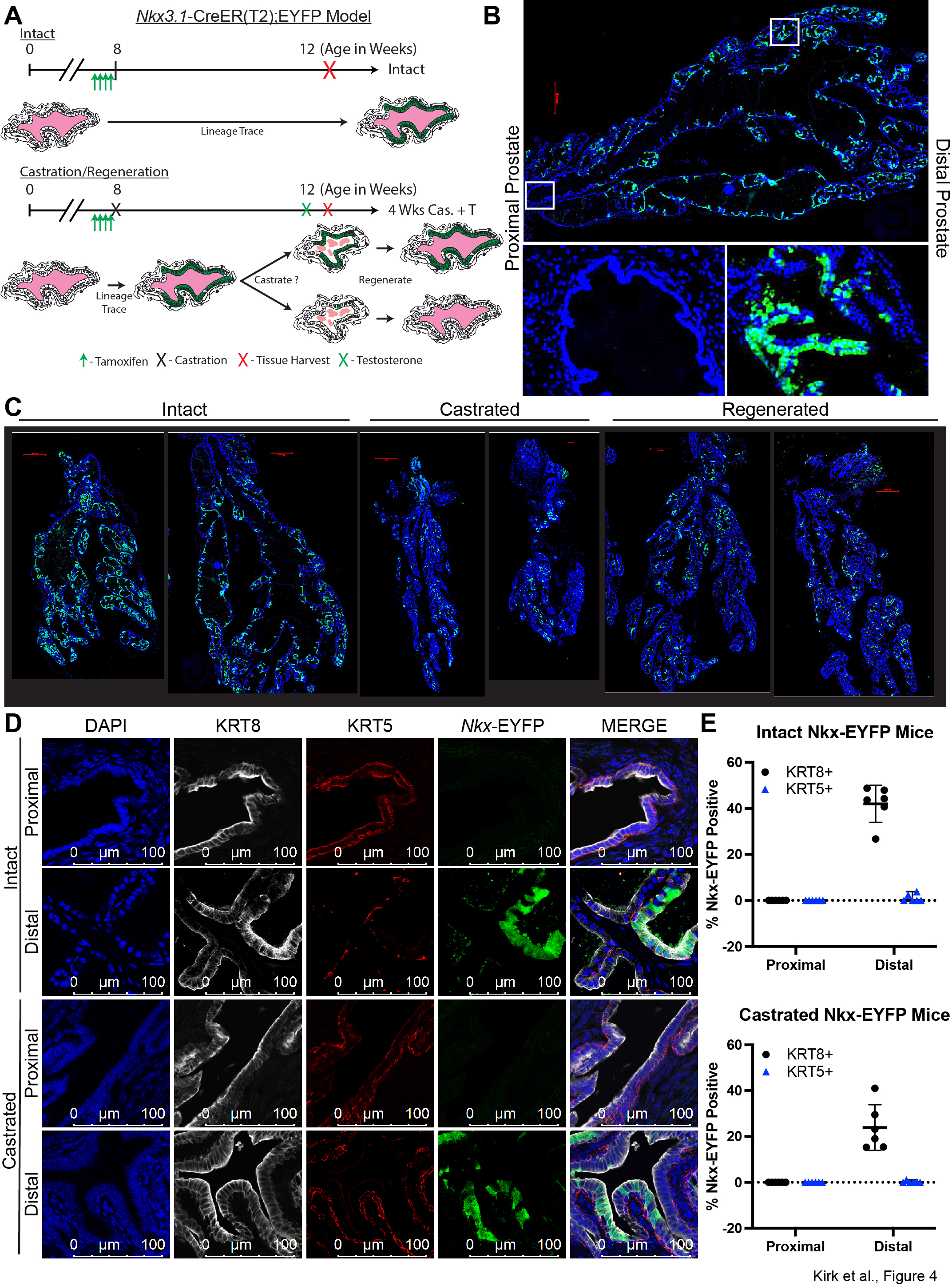
*Nkx3.1-*CreER(T2) lineage tracing demonstrates that L7 can give rise to L1. **(A)** Schematic demonstrating expected outcomes of lineage-tracing studies when EYFP tracer is activated by Tamoxifen in intact and castration/regeneration settings. **(B)** In intact prostate, lineage tracing labeled mainly distal luminal cells (Green) with sparse labeling in proximal region. White boxed regions are enlarged (below). **(C)** Representative whole-mount images of pre-castration labeled *Nkx*;EYFP APs from intact, castrated, and regenerated prostates. **(D)** Representative images of pre-castration labeled *Nkx*;EYFP AP double stained for KRT8 (AF-647) or KRT5 (AF-594) in proximal and distal regions of intact and castrated prostates. **(E)** Quantification of proximally and distally labeled EYFP^+^ luminal and basal cells from intact and castrated prostates. Scatter plots represent the mean ± SD of 3 independent regions of interest (ROI) in proximal and distal AP from 2 independently labeled mice.

The above results demonstrated that pre-existent *Clu^+^* cells (L5) *do not* give rise to castration-resistant L1 cells and suggested that the L1 population might have derived from distally located and differentiated luminal cells. To test this possibility, we employed the *Nkx3.1* driven Cre-ER(T2)*;EYFP* (*Nkx*;EYFP) lineage tracing model^38^. Labeling of *Nkx;*EYFP model prior to castration and following tagged cells through one round of castration/regeneration will demonstrate either a massive loss of EYFP-labeled cells or survival of this population upon castration/regeneration (Figure 4A). As previously reported^38^, in intact AP the EYFP^+^ cells were found predominantly in distally located luminal cells with little proximal labeling (Figure 4B). Examining multiple APs from intact, castrated, and regenerated mice revealed that distally labeled *Nkx;*EYFP*^+^* luminal cells survived castration and, importantly, gave rise to regenerated prostates (Figures 4C, S6C). Quantification of EYFP^+^ KRT8 and KRT5 expressing cells in intact and castrated conditions corroborated these findings (Figures 4D-E). Examination of the *Nkx*:EYFP model in post-castration setting has been done extensively^38^ and hence not repeated herein. These results demonstrated that differentiated *Nkx3.1-*expressing cells can survive castration and become part of the *Clu*^+^ L1 population, since L1 is the *only* distally located population in AP post-castration (Figures 1B, 2B).

Lastly, we utilized a *Krt5-*CreER(T2);tdTomato (*Krt5*;Tomato) lineage tracing model^39, 40^ to address whether basal (stem/progenitor) cells could give rise to L1 during castration. In adult mouse prostate, although basal and luminal compartments are independently maintained by their own progenitor cells^22, 23, 29, 30, 41^, basal progenitor cells could potentially generate luminal cells during castration and can differentiate into luminal cells upon luminal cell loss by anoikis^41^. Our results showed that *Krt5;*Tomato mice exclusively labeled KRT5^+^ basal (but not KRT8^+^ luminal) cells in both proximal and distal regions of intact, castrated, and regenerated prostates (Figures S6D-E). Therefore, the castration-resistant L1 population was not derived from basal cells.

Our results indicate that castration-resistant L1 cells *do not* arise from proximally or distally located L5 nor from basal cells. Instead, L1 *can* arise from distally located *Nkx3.1*-expressing L7/L10 cells, suggesting emergence of L1 as a result of epigenetic plasticity in differentiated luminal cells.

### Chromatin and differentiation trajectory analysis reveal acquired stemness in L1 cells

We employed CytoTRACE, a program designed to predict developmental potential or stemness^42^ between cells of similar lineage to further elucidate the relationship between luminal populations (Figure 5). CytoTRACE revealed high stemness (Score closer to 1) in L5 and, surprisingly, in L1 and L3 as well in both independent single-cell sequencing projects (Figure 5A). Differentiated luminal cells L2, L4, L6, L7, L8, and L10 all had low stemness scores, although there was some diversity within these populations. L9 was found to have an intermediate score, further supporting the transitory nature of this population. These results confirmed the stemness associated with the proximal niche (L5), but also interestingly demonstrated that castration induced increased stemness in distal luminal cells (L1/L3).

**Figure 5.**
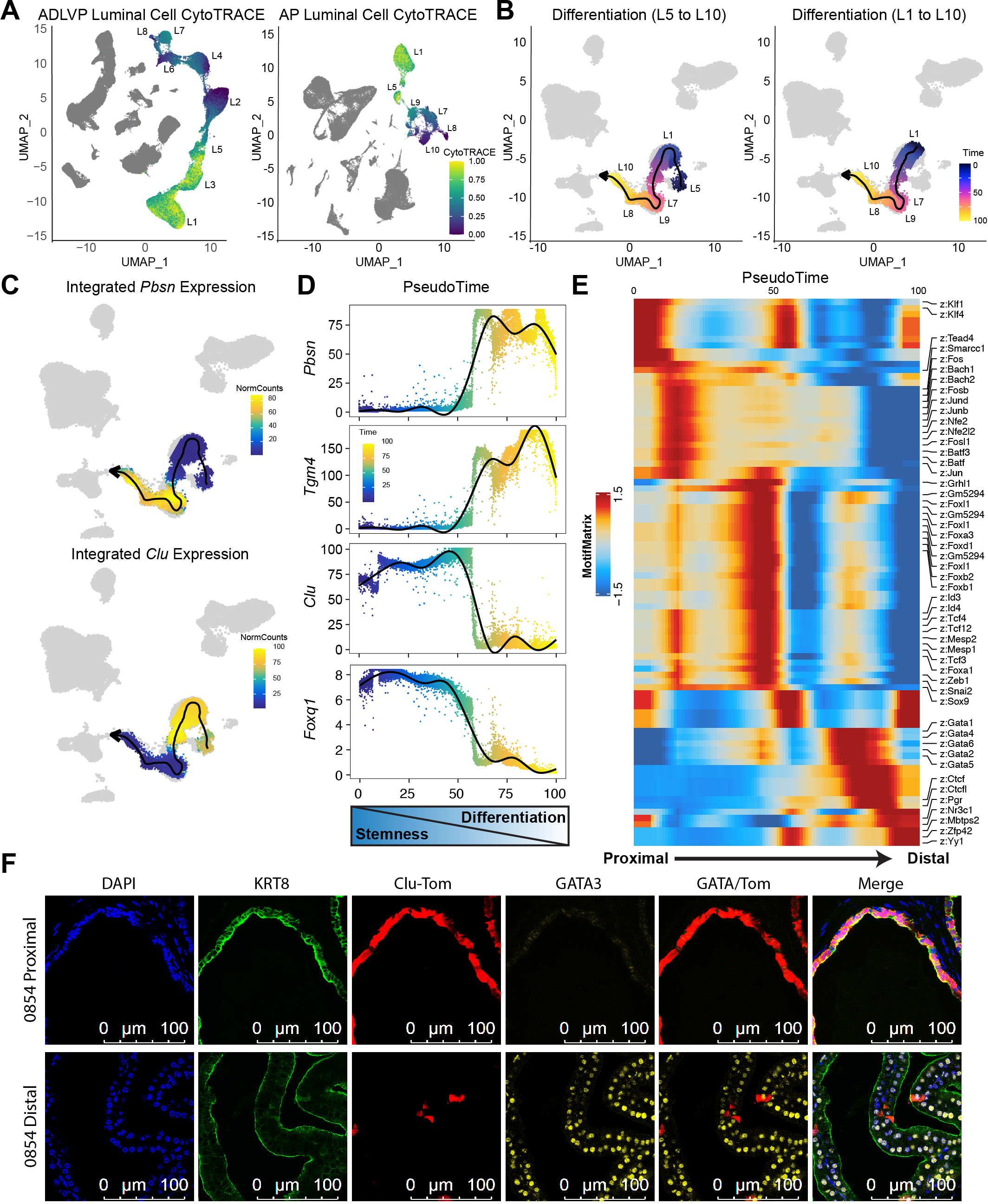
Chromatin analysis identifies dynamic changes along a differentiation trajectory. **(A)** CytoTRACE analysis of luminal cells from ADLVP and AP scRNAseq demonstrates a higher stemness score (closer to 1) in castration-resistant clusters L5/L1/L3 relative to differentiated luminal clusters L2/L4/L6/L7/L8. **(B)** Differentiation trajectory lines starting from L5 or L1. Time progresses from 0 (blue) to 100 (yellow) along the trajectory lines. **(C)** Integrated RNA expression of select markers along the luminal cell differentiation trajectory overlayed on scATACseq UMAP. **(D)** Normalized integrated RNA read counts for individual cells plotted against the differentiation timeline demonstrates an increase in differentiation markers over time. **(E)** Heatmap representation of the changes in motif enrichment along the L5 to L10 differentiation trajectory. **(F)** IF staining of GATA3 supports the differential regulation of GATA motifs observed between proximal stem-like cells and distally located differentiated luminal cells.

We utilized the stemness relationships of luminal cell populations to construct a pseudotime trajectory along the differentiation path (Figure 5B). Analyzing changes along these trajectory lines revealed changes in relation to developmental timelines with L5 as the root, or along castration-induced timelines with L1 as the root. Using the integrated dataset, expression changes along the L5 pseudotime demonstrated increased expression of *Pbsn* and *Tgm4* (Figures 5C-D). Conversely, *Clu* and *Foxq1* were found to be highly expressed in undifferentiated stem-like populations. Analysis of motif enrichment demonstrated a clear differentiation path for prostate luminal cells (Figure 5E). Transitioning from the proximal niche to distal prostate, we observed that *Klf* family motif-regulated stem cells activated a *Fox* driven program, and that reaching full differentiation potential required *Gata* engagement followed by direct *Ar* motif-mediated transcriptional output (Figure 5E). Confocal analysis revealed higher GATA3 expression in *Clu*-negative distal luminal cells relative to *Clu-*positive proximal niche (Figure 5F), confirming late activation of GATA3 during differentiation.

### Pathway analysis and therapeutic studies corroborate JUN/FOS dysregulation during castration resistance

To better define molecular changes associated with castration resistance, differentially expressed genes (DEGs) were calculated between intact L7/L10 and castration-resistant L1 as well as between L5 and L1 (Table S6). There were 1,510 DEGs identified between L1 and L7/L10 populations. Ingenuity Pathway Analysis^43^ (IPA) revealed that L1 was enriched in integrin, unfolded protein response, MSP-RON, PI3K/AKT, HER-2, and JAK/STAT signaling pathways, among others (Figure 6A; Table S6). Relative to L5, L1 had 728 DEGs that were uniquely involved in NRF2-mediated oxidative stress response, glutathione-mediated detoxification and HIF1*α*, aryl hydrocarbon receptor and p53 signaling (Figure 6B; Table S6). Strikingly, IPA Upstream Regulator analysis revealed many ‘deregulated’ TFs in L1 that were also predicted to be deregulated by scATACseq, including JUN/FOS, SMAD3/4, CTNNB1, TCF7L2, RELA/NF*κ*B1, and STAT3 (Figure 6C). Drugs predicted to be deregulated (i.e., drugs that could mimic the observed changes in DEGs) included, among others, doxorubicin, palbociclib, finasteride, androgen, LY294002 and imatinib (Figure 6D). Of interest, finasteride would simulate a castration response in prostate and was predicted to be activated in L1; conversely, androgen would stimulate regeneration and was predicted to be deactivated in L1 (Figure 6D). Lastly, we performed gene set enrichment analysis (GSEA) using the hallmark gene sets in MsigDB, and results showed that L1, relative to L7/L10, was enriched in TNF*α* signaling via NF-*κ*B, epithelial to mesenchymal transition, IL2-STAT5 signaling, TGF*β* signaling, inflammatory response, and KRAS signaling_UP (Figure 6E-F). Cumulatively, IPA and GSEA demonstrated that several pathways, including JUN/FOS, NF-*κ*B, PI3K/AKT, ZEB1, WNT/*β*-catenin, EGFR/ERBB and JAK/STAT signaling were all contributing to castration response in L1. Predominantly located to the stromal layer in intact prostates, the NF-*κ*B regulatory unit RelA was highly upregulated in castration-resistant L1 (Figure 6G). Furthermore, upregulated genes in L1 may be directly mediating these pathways, such as *Alox12* mediating inflammatory response, *Btc* EGFR/ERBB signaling, and *Lgr4* WNT/*β*-catenin signaling (Figures 6H, S3F). Given the upregulation/dysregulation of these pathways, the prostate regression and regeneration responses might be impacted upon their perturbation. Therefore, castrated mice were treated with inhibitors to identified pathways (dose and schedule in Figure 6I). HDAC (Panobinostat) and WNT (XAV-939) inhibitors were found to impair prostate castration survival/regeneration, while Bortezomib (proteosome/NF-*κ*B inhibitor) and Capivasertib (AKT inhibitor) both slightly inhibited prostate survival/regeneration although not to a significant degree (Figure 6J). Surprisingly, ML355 (Alox12 inhibitor) and Tanzisertib (JNK/AP1 inhibitor) were both found to be protective of prostate castration/regeneration (Figure 6J), suggesting that these pathways may play a more complex role in regulating castration response.

**Figure 6.**
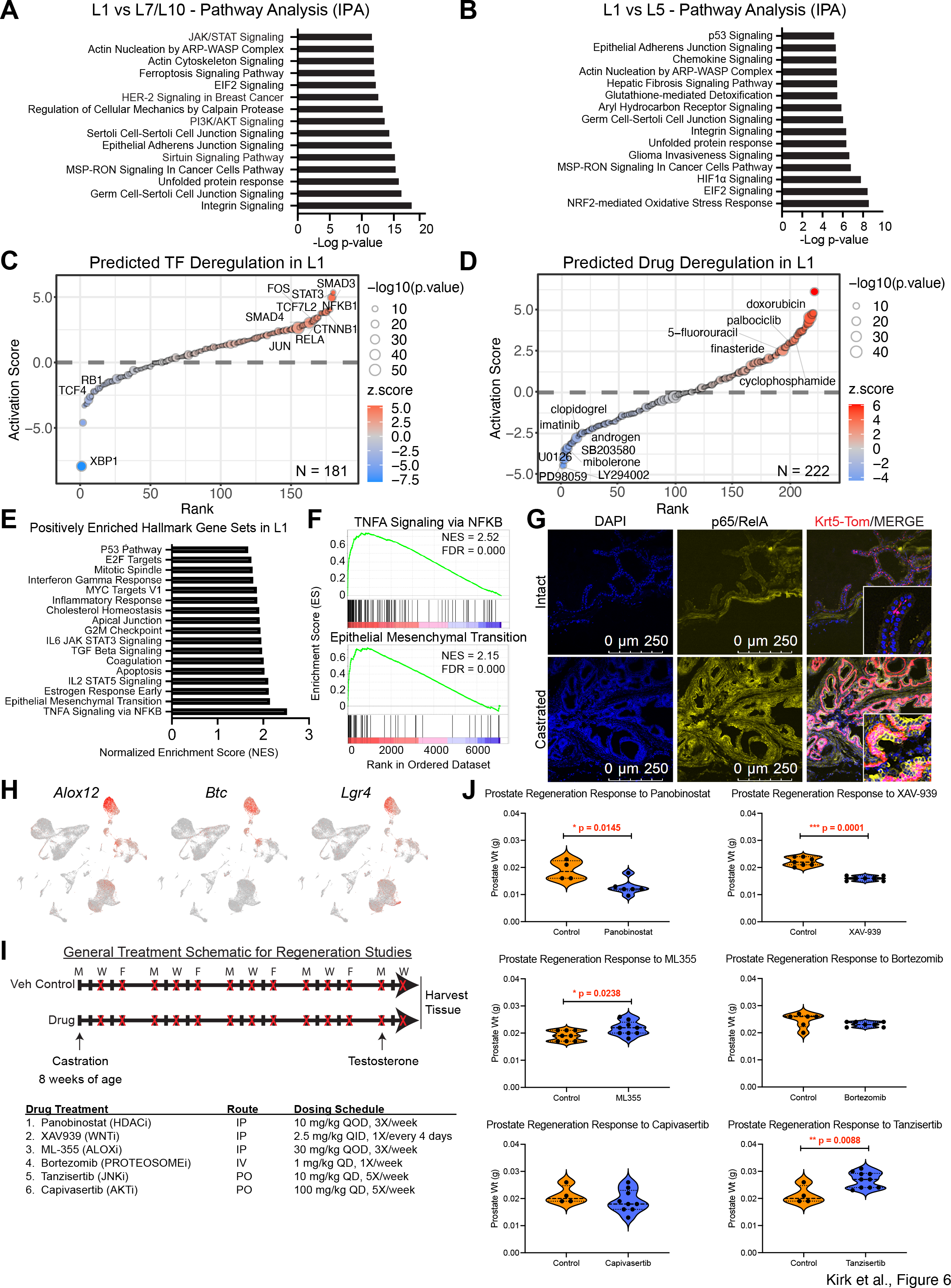
Pathway analysis reveals regulatory mechanisms driving L1 castration resistance. **(A-B)** IPA of DEGs between L1 and L7/L10 (pooled) populations (A) or between L1 and L5 (B). The top 15 deregulated pathways ranked by -Log(p-value) are displayed. (**C-D**) IPA prediction of top deregulated TFs (C) and chemical drugs (D) in L1 based on curated datasets. Predicted deregulations are presented as rank-ordered plots of the predicted activation z-score for each factor. Bubble point size indicates -Log10(p-value). (E) Hallmark gene set enrichment analysis (GSEA) of DEGs in L1 relative to L7/L10 identifies positively enriched gene sets. (F) Rank-ordered plots of two positively enriched gene sets in L1. (G) Confocal images of p65/RelA staining in intact and castrated *Krt5*;Tomato lineage-traced APs. (H) Representative UMAP overlays for select markers deregulated in pathway analysis. (I) Treatment schema using inhibitors of pathways deregulated in L1 with castration. Drug name, target, route of administration and dosing schedule are indicated. (J) Prostate weight of mice undergoing castration/regeneration treated with indicated drugs or vehicle.

### L1-like cells are present in human PCa samples treated with ADT or ARSIs

To determine the human prostate and PCa relevance, the AUCell function from SCENIC^32^ was again applied to specific gene sets to determine their relative activity within individual cells of the AP dataset. Specifically, signatures associated with PCa response to ADT/ARSI^44–46^ were analyzed. We observed that genes downregulated by ADT/ARSI were predominantly enriched in differentiated luminal populations (Figure 7A, Down; Table S7) whereas genes upregulated by ADT/ARSI were found to be enriched mainly in L1/L5 and basal cells (Figure 7A, Up). These latter observations suggested that castration-resistant human PCa cells may be both L1 and basal-like in nature. In support, using advanced PCa signatures^47^, we found that the PCa squamous cell signature (Squam, KRT6A^+^) was enriched in both basal and L1 cells (Figure 7B). Interestingly, the amphicrine (AR^+^/NE^+^) signature was enriched in differentiated luminal and stromal cells (Figure 7B; Table S7), possibly suggesting more phenotypic plasticity in these tumors. The NE signature was enriched in the few NE cells identified, and finally the double-negative (AR^-^NE^-^) tumor signature was generally enriched everywhere except AR^+^ differentiated luminal cells (Figure 7B).

**Figure 7.**
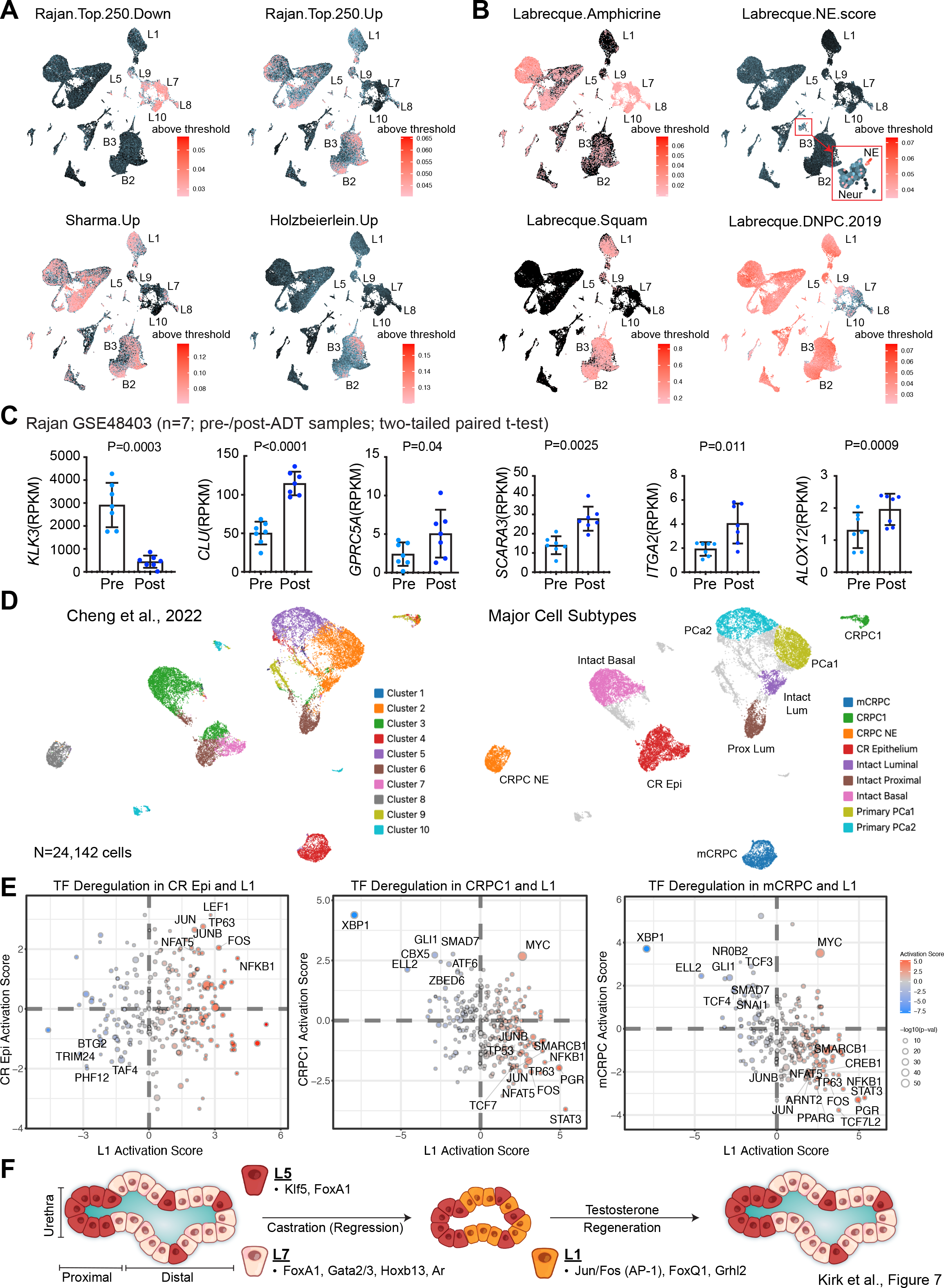
Relevance of L1 to human PCa response to castration. **(A)** AUCell overlays of gene expression signatures associated with neo-adjuvant ADT response in primary PCa patients from Rajan et al.^44^ (GSE48403), Sharma et al.^46^, and Holzbeierlein et al.^45^ Down/Up, gene signatures lost or gained, respectively, with chemical castration. **(B)** AUCell enrichment of gene signatures related to CRPC subtypes in Labrecque et al.^47^. **(C)** Representative L1 genes upregulated in post-ADT PCa in the Rajan dataset. **(D)** Re-analysis of the Cheng et al.^48^ dataset examining human PCa response to castration in 24,142 single cells identified 10 cell clusters and 9 PCa cell subtypes. **(E)** Scatter plots of IPA-predicted TF deregulation in CR Epi, CRPC1, and mCRPC cell subtypes compared to predicted L1 TF deregulation. Activation score color and -Log10(p-value) bubble size are indicative of the L1 population. DEGs from human subtypes used in TF analysis were generated relative to the Intact Luminal subtype. **(F)** Schematic depicting the major differences in proximal L5 stem/progenitor cells, differentiated L7 cells and reprogrammed castration-resistant L1 cell population (see Text).

In addition to examining gene signature enrichment (Figure 7A-B), we also analyzed the expression of candidate genes upregulated in L1 in several neoadjuvant ADT datasets^44, 46^. While the AR target genes *KLK2/KLK3* were downregulated with ADT as expected, the L1-specific genes *CLU*, *GPRC5A*, *SCARA3*, *ITGA2*, and *ALOX12* were upregulated (Figures 7C, S7A). In contrast, the L5- specific genes *PSCA*, *WFDC2*, *SPINK5*, and *CYP2F1* remained unchanged with ADT (Figure S7B-C), suggesting that, consistent with our observations in castrated mouse prostate, the L5-like cells in human PCa were not expanded with castration. IPA Upstream Regulator analysis of DEGs associated with ADT^44^ revealed that the predicted TF and Drug deregulations were highly concordant with the changes found in castration-resistant L1 (Figure S7D,E). Specifically, predicted activation of JUNB, CTNNB1/TCF7, and STAT3 pathways was all observed. These findings further support that the biology associated with castration-resistant L1 is directly relevant to human PCa ADT response.

To determine whether our ADLVP and AP single-cell studies could be corroborated at the single-cell level in human PCa, we re-analyzed a recently published single-cell study^48^ containing matched primary PCa and benign samples, CRPC, and NEPC (Figures 7D-E, S7F-I). 24,142 high-quality cells from 11 samples (Figure S7F) were clustered using a K-means 10 algorithm to identify 10 main cell clusters (Figure 7D). Using the sample annotations, the original publication, and lineage-specific markers, we designated 9 major cell populations or subtypes (Figures 7D, S7F,G). Using a recently reported 6-gene panel^49^ that included *AMACR*, we were able to identify and segregate normal from cancerous cells (Figure S7G). Since most of these PCa samples came directly from the prostate, it is not surprising to see a percentage of normal cells, even within the CRPC collected from the primary gland. After stratifying normal from cancer cells, we identified two types of primary PCa (PCa1 and 2), three types of CRPC (one metastatic (mCRPC) and two in the prostate (CRPC1 and CRPC-NE)), and four normal cell populations (intact luminal, intact proximal, intact basal and castration-resistant (CR) epithelium) (Figure 7D). Candidate and globally distinguishing genes were found for each major cluster/population (Figures S7H,I): proximal luminal cells were identified by *PSCA* expression, basal cells by *KRT5/14*, AR signaling active luminal cells by *KLK3*, primary PCa by *ERG*, and CR Epithelium by *CLU*, *ANXA10*, and *FABP4*.

Next, DEGs between intact luminal and the non-NE CRPC clusters were calculated (Table S7). DEGs were again assessed by IPA Upstream Regulator analysis and compared to the deregulations observed in L1. Strikingly, several TF deregulations predicted for normal CR Epi positively corresponded to those of the L1 phenotype, namely, JUN/FOS, LEF1 and NF*κ*B (Figure 7E). Furthermore, TF deregulation in CRPC1 and mCRPC was inversely associated with TF deregulation observed in L1 (Figure 7E), suggesting reactivation of androgen-regulated pathways for these CRPCs as supported by *KLK3* expression (Figure S7H) and the well-established CRPC mechanisms. Interestingly, L1-specific TFs, JUN/FOS, STAT3, WNT and NF*κ*B, were predicted to be deactivated with CRPC emergence, possibly suggesting AR-mediated suppression of these pathways. Altogether, these findings highlight the relevance of our ADLVP and AP datasets in understanding human PCa response and resistance to ADT/ARSIs.

## DISCUSSION

mCRPC is estimated to kill >34,000 American men in 2023^10^. This high mortality, despite clinically effective ARSIs such as enzalutamide, is associated with our fragmented understanding of the evolutionary mechanisms and cell(s)-of-origin of CRPC. Our current study, by combining multi-omics single-cell approach and lineage tracing in the mouse prostate in response to castration and regeneration, greatly advances our knowledge in both fronts. *First*, we have generated a robust single-cell mouse prostate atlas consisting of 229,794 cells and identified novel cell populations and response-specific markers in intact, castrated and regenerated prostates. Our robust single-cell atlas also allows refined stratification of previous single-cell datasets and serves as an invaluable resource for future studies. *Second*, we report that castration induces dramatic cell population changes in all 3 major prostate cell lineages, i.e., luminal, basal, and stromal. Importantly, these dynamic cellular shifts are supported at both the transcriptomic and chromatin levels. *Third*, of relevance to understanding the cell(s)-of-origin of CRPC, independent lineage tracing of proximal and distal luminal cell populations shows that the major castration-surviving luminal population, L1, does not arise from the expansion of pre-existent *Clu*-positive luminal progenitors nor from basal cells. Instead, castration-resistant luminal cells are derived from the NKX3.1^+^ differentiated luminal population. *Fourth*, global chromatin accessibility analysis reveals that, although castration-reprogrammed luminal cells manifest a stemness score equivalent to that of the proximal luminal stem/progenitor (L5) cells, these two cell populations possess distinct chromatin landscapes and TF occupancy profiles. Specifically, the proximal luminal niche is predominantly regulated by *Klf1/5* and *Fox* regulatory motifs whereas *Jun/Fos, Fox, Grhl,* and *Gata* motifs characterize the castration-resistant luminal cells in the distal prostate (Figure 7F). *Fifth*, pathway analysis unravels deregulation of JUN/FOS, PI3K/AKT, eicosanoid, WNT/*β*-catenin, NF*κ*B and JAK/STAT signaling in luminal cells responding to castration, and therapeutic targeting of some deregulated pathways disrupts plasticity-induced castration resistance. *Finally*, interrogation of human PCa datasets uncovers direct relevance of our results in the mouse prostate to human PCa castration response and also reveals that mouse and human prostate luminal cells activate many of the same signaling pathways during evolution to castration resistance.

Single-cell analysis in high cell numbers (Figure S2C) has allowed us to identify novel luminal, basal and stromal cell populations in both intact and castrated prostate. For example, there are 6 differentiated luminal cell populations (L2, L4, L6, L7, L8 and L10) in intact prostate (excluding L5) and castration reduces this heterogeneity to 2 populations (L1/L3). Castration similarly reduces cellular heterogeneity in stromal and basal compartments, generating castration-resistant Me1 and B2 populations. AR signaling is known to play an important role in regulating stromal cell functions and stromal-epithelial crosstalk in the prostate^50, 51^, so it is not surprising to see changes in mesenchymal cells nor the high level of *AR* motif enrichment in mesenchymal populations. The Me1 population that survives castration loses differentiation markers *Col3a1, Col1a2* and *Gsn*, suggesting the critical role of AR signaling in maintaining stromal cell differentiation. Of interest, the Me1 cells also gain expression of *Apod*, which has been reported to mark the peri-epithelial (APOD^+^) fibroblasts^7^. Traditionally, the role that AR plays in basal cells is thought to be negligible since AR signaling is not required for basal cell survival^15, 16^. However, it was recently demonstrated that *AR* deletion in basal cells did impact their ability to survive and differentiate into luminal cells^52^. Intriguingly, castration-resistant B2 cells upregulate, among other genes, *Krt5,* suggesting castration-induced keratinization in basal cells. In support, human PCa subjected to short-term ADT (Figure 7A) and squamous subtype of CRPC (Figure 7B) also exhibit enhanced basal-cell gene expression. Altogether, our results demonstrate that castration causes profound population changes in not only luminal cells but also stromal and basal cells.

Studying castration/regeneration responses in mouse prostate has great implications in understanding human PCa response to ADT/ARSIs and potential cell(s)-of-origin of CRPC. In both mouse and human prostate, the urethra-proximal prostate harbors quiescent epithelial stem/progenitor cells that are *intrinsically* refractory to castration^1, 3, 5, 6, 25–27, 53–56^. The proximal epithelial cells under-express differentiation regulators AR and NKX3.1 but overexpress stem cell molecules such as Sca-1, SOX2 and RUNX1^53, 54, 57, 58^; this study). Due to their intrinsically castration-resistant nature, the proximal progenitor cells have been thought of as the cells that would survive castration and mediate regeneration/recurrence. Further, a recent study suggests that castration reprograms differentiated luminal cells to become proximal-like and castration-resistant^8^. The proximal progenitor cells reported in these earlier studies correspond to our L5 cells, which express a unique repertoire of phenotypic markers and TFs (Figure 7F). L5 cells are also scattered across distal ducts and acini. Interestingly, the distally located L5 cells have also recently been shown to expand during castration/regeneration to give rise to regenerated prostates^4^. However, our studies herein indicate that 1) the L5 cell population, both proximally and distally located, does not expand in response to castration; 2) the major castration- reprogrammed luminal cells, L1, do not become (proximal) L5-like during castration; 3) L1 and L5 represent two distinct populations; and 4) L5 cells do not give rise to L1 cells during regeneration and L1 cells can regenerate the prostate upon testosterone re-administration.

The two independent scRNAseq experiments in >180,000 cells identified 3 castration-resistant luminal cell populations, the pre-existent L5 and *de novo* L1 and L3. By focusing on L1, we provide multiple lines of evidence that dissect the relationship between L1 and L5 and distinguish L1 from L5. *First*, scRNAseq shows that the L5 in AP does not expand during castration (Figures 1B, 2C). *Second*, they possess distinct gene profiles and express different marker genes – although the two populations share a set of genes associated with stem cells and castration resistance like *Clu, Krt19, Foxq1,* and *Tacstd2* (Figure S2F), L1 cells do not express L5-unique genes such as *Psca* and *Wfdc2*. Instead, L1 cells uniquely express *Alox12, Btc, Upk3a, Prr9, Itga2* and *Krt6a* (Figures S2G, S3F). *Third*, remapping Karthaus dataset using our reference atlas also reveals L1 and L5 to be unique populations. *Fourth*, lineage tracing using the *Clu;*Tomato model shows that proximal L5 does not expand into the distal prostate and pre-existent scattered L5 population does not expand in the distal region upon castration. In contrast, lineage tracing using the *Nkx3-1*;EYFP model demonstrates that differentiated luminal cells can give rise to L1. We speculate that only a subset of *Nkx3-1^+^* cells have the ability to ‘de-differentiate’ into L1 cells upon castration and a candidate subset is L7. In intact AP, although L10 is a differentiated luminal cell population similar to L7, only L7 expresses some L1 markers such as *Sox9, Cxadr* and *Itga6*. Also, *Nkx3-1* is expressed in a higher percentage of L10 than L7 cells, 55% vs 39%, and lineage-traced cells in the *Nkx3-1*;EYFP model, although maintained with castration, are reduced relative to intact mice, suggesting that L10 might be more sensitive to castration than L7. *Fifth*, L1 and L5 cell populations are enriched in different signaling pathways (Figure 6B). *Finally*, our scATACseq data reveal distinct chromatin landscapes in L1 vs. L5 cell populations.

The scATACseq analysis corroborates key findings from the scRNAseq showing L1, B2, and Me1 emergence with castration. scATACseq also reveals the dynamic nature of the chromatin landscape after castration/regeneration, with L1 shifting back toward the more differentiated landscape of L7 within 96 hr of testosterone treatment. Regenerated L9 was virtually indistinguishable from differentiated L7 at the chromatin level. Similarly, castrated B2 and Me1 were rapidly restored to intact chromatin landscapes upon regeneration. Interestingly, scATACseq identified 2 unique populations not identified by scRNAseq, specifically a basal intermediate (BInt) and a L1 intermediate (L1Int). L1Int integrated with L1 but seems to be an intermediate population between L1 and L7/L9. BInt integrated with B3/B2 but seems to serve as a castration-insensitive root for both populations. How these populations contribute to the overall biology of the prostate and castration response awaits further studies.

Critically, our scATACseq study identifies known TF such as FOXA1^59, 60^, HOXB13^61, 62^ and GATA2/3^63, 64^ and also nominates other TF including JUN/FOS (AP1), KLF and GRHL in differentially regulating differentiated luminal, pre-existent L5 and castration-reprogrammed L1 cells (Figure 7F). FOXA1, a pioneering TF, acts upstream of AR and co-localizes with AR greater than 50% of the time^65^. We find that although *Fox* motifs are enriched in L5, L1, L7, and L10 populations, there is diverse *Foxa1* expression across luminal cell subtypes and interestingly, the castration-resistant L1/L3/L5 have higher expression of *Foxq1* (Figures 1E, S3F). Unlike the *Fox* motifs, both *Hox* and *Gata* family motifs are most highly enriched in intact L7/L10 luminal cells but not L1/L5 populations (Figure 7F), supporting that HOX and GATA members normally serve as AR co-regulators in promoting luminal differentiation. Interestingly, both *Grhl2* expression and *Grhl2* motif are increased in L1 cells. It is unlikely that GRHL2 is serving as an AR co-activator^66^ in this castrated context. GRHL2 has been shown to collaborate with FOXA1 to drive tamoxifen resistance in breast cancer^67^; thus it’s tempting to speculate that GRHL2 may be serving a similar role in regulating castration resistance in prostate. The proximal L5 is uniquely enriched in *Klf* family motifs (Figure 7F), which have been reported to play a role in regulating proximal stem cells of the prostate^68^. Other factors including SOX2^54, 69^ and RUNX1^58^ have also been implicated in regulating the proximal progenitor niche; however our data does not support a large role for these molecules relative to the KLF family. Consistently, several stem cell genes associated with castration resistance, including *Bmi1, Cdh2, Sox2, Zeb1* and *Runx2*, are not increased in castration-resistant L1/L3/L5 cells (Figure S2I). Overall, examining regulatory factors along a stemness timeline, we find that KLF and FOX family members regulate the primitive stem cell niche of the proximal gland, but as cells differentiate in response to androgen, they become enriched in GATA, HOX, and AR regulatory motifs, with AR being the last to emerge (Figure 7F).

Strikingly, the *Jun/Fos* (AP-1) regulatory motifs are identified as the top enriched motifs in castration-reprogrammed L1 cells. AP-1 has been implicated in many cancers including PCa^70, 71^. c-JUN was reported to be an AR co-activator^72, 73^ whereas c-FOS was antagonistic^74^. It was later shown that AP-1 activation by TPA antagonized AR activity^75^ and that AR repressed c-FOS expression^76^. An interesting recent study reports that various combinations of AP-1 family members regulate the level of differentiation and cellular plasticity in melanoma^77^, prompting us to speculate on potentially antagonistic roles of c-JUN and c-FOS in regulation AR signaling with c-JUN being pro-differentiation (AR-active) and c-FOS important for plasticity and stemness (AR-inactive). *De novo* motif analysis from the scATACseq shows the presence of AP-1 motifs in both differentiated L7 and castrated L1. Under intact conditions, c-JUN may be serving as a co-activator of AR signaling^72, 73^, functioning similarly to GATA2 in regulating AR activity^64^. During castration, *c-Fos* expression is upregulated in L1 relative to L7/L10 (Table S6), likely due to loss of AR repression^76^. The increased *c-Fos* expression may shift AP- 1 composition to a JUN/FOS heterodimer, and thereby alter chromatin organization and differentiation state of luminal cells, as suggested by the studies from Comandante-Lou et al.^77^

Pathway analysis has identified integrin, eicosanoid, PI3K/AKT, HER-2, and JAK/STAT signaling to be deregulated in castration-resistant L1, and Upstream Regulator analysis has implicated JUN/FOS, STAT3, SMAD3/4, NF*κ*B, and CTNNB1 TFs most likely to be deregulated with castration. JUN/FOS deregulation is fully consistent with our scATACseq analysis and Tanzisertib (CC-930), a JNK/AP1 inhibitor, surprisingly, promoted prostate regeneration (Figure 6J). Similarly, ML355, an inhibitor of eicosanoid-metabolizing enzyme Alox12, which is specifically and significantly upregulated in L1 cells (Figures S2G, S3F), also promoted prostate regeneration. These unexpected findings suggest that castration-induced AP1 activation and upregulation of A12lox-mediated eicosanoid metabolism may functionally inhibit the regenerative capacity of L1 cells. On the other hand, inhibitors of WNT/*β*-catenin (XAV-939), AKT (Capiversatib), HDAC (Panobinostat) and proteosome/NF*κ*B (Bortezomib) all demonstrated an inhibitory effect or trend on prostate regeneration (Figure 6J), implicating these pathways in driving L1 generation and L1 regenerative activities. We also find JAK/STAT, which has received a lot of attention recently in relation to advanced PCa lineage plasticity^78–80^, to be one of the pathways deregulated in L1; however, we do not see any deregulation or enrichment of *Stat* motifs with castration in our scATAC studies. Furthermore, STAT chromatin recruitment has not been demonstrated in any of the recent studies, leaving us to wonder how JAK/STAT may be regulating treatment-induced plasticity.

Our results in the mouse prostate can be directly linked to human PCa response and resistance to ADT/ARSIs. For example, the L1 signature, compared to the differentiated L7/L10 gene signatures, is enriched in post-ADT PCa and in aggressive CRPC subtypes such as double-negative and squamous CRPC. The association of L1 signature to squamous CRPC is also supported by remarkable upregulation of numerous *Krt* genes, including *Krt4, Krt6a, Krt8, Krt18*, and *Krt19* and ‘general’ epithelial genes such as *Cdh1, Epcam* and *CD24a* in castration-resistant L1/L3/L5 cells (Figure S2D,F), again suggesting, as in basal cells, castration-induced epithelial keratinization as an underlying mechanism of castration resistance in luminal cells. Significantly, many L1/L5 or L1-specific genes are upregulated in post-ADT human PCa; in contrast, the L5-unique genes remain unchanged in post-ADT PCa, supporting that the proximal L5 cell population is also not expanded in human PCa undergoing ADT. Remapping a recent human PCa scRNAseq study^48^ reveals that pathways deregulated in mouse L1 are also deregulated in castration-resistant human prostate luminal cells, and, interestingly, these pathways are inversely correlated in CRPC cell populations, consistent with re-activation of AR signaling in end-stage CRPC. Furthermore, analysis of human datasets suggests that JUN/FOS deregulation may play a key role in developing resistance early in response to neoadjuvant ADT (Rajan dataset), but eventually is overcome in CRPC (Cheng dataset). Recent ATACseq in organoid and PDX models identified 4 main subtypes of advanced CRPC, AR-dependent, NEPC, WNT-dependent, and stem cell-like (SCL)^81^. Remarkably, CRPC-AR is found to be regulated by *AR, FOX, GATA* and *HOX* family motifs, matching our L7 population, and CRPC-WNT by *TCF, KLF, SOX* and *RUNX* family motifs, similar to our L5 population. Most notably, CRPC-SCL is regulated by *JUN/FOS, NFκB*, and *TP63* family motifs, closely matching the L1 population. In addition to these findings, a study examining the chromatin accessibility landscape of human cancers identified 7 distinct accessibility clusters in PCa patients^82^. These clusters, identified from bulk ATACseq samples, are enriched in motif combinations of *AR/HOX, FOX, FOX/KLF, AR/GATA*, and *JUN/FOS/TP63* family members, suggesting that the clinically diverse subtypes of PCa and CRPC are regulated by different transcriptional programs operating in differentiated (L7), castrated (L1) or even primitive progenitor (L5) populations. In summary, our mouse studies offer clear clinical relevance for understanding castration resistance and CRPC.

## CONCLUSIONS

The work presented here offers a framework for understanding prostate luminal cell response to castration. Integrated scRNAseq and scATACseq in ∼230,000 mouse prostate cells have allowed us to thoroughly delineate the heterogeneity, dynamic changes and epigenetic basis of cell populations during castration and regeneration. The major castration-resistant luminal cell population, L1, is highly relevant to ADT/ARSI treated human PCa. The deciphered L1 cell origin, i.e., being reprogrammed by castration from differentiated *Nkx3.1*-expressing luminal cells suggests that in clinically treated PCa, in addition to phenotypically undifferentiated (L5-like) cancer stem cells^48, 81, 83, 84^, differentiated PCa cells may also serve as the cells-of-origin for CRPC as a consequence of treatment-induced epigenetic plasticity. We identify JUN/FOS (AP1) as one of the major TFs driving castration-induced L1 reprogramming. Finally, we offer several potential therapeutic options to curb the treatment-induced plasticity such as combining ADT/ARSIs with HDAC, WNT/*β*-catenin or AKT inhibitors.

## STUDY LIMITATIONS

We identified two *de novo* castration-resistant luminal populations, L1 and L3, but the functions, origin and human relevance of DLVP-derived L3 need to be fully characterized in future studies, so as the castration-resistant Me1 and B2 populations. We also identified rare luminal cell subtypes in intact mice such as L8, which express *Lars2, Tcaf3,* and *Gm42418*, but their functional relevance to the prostate remains to be elucidated. It will be very interesting to dissect exactly which subset of Nkx3.1^+^ cells give rise to L1 in response to castration. Further chromatin and population-specific studies are warranted to confirm the lineage-restricted regulation of various TF programs. Moreover, the contribution of JUN/FOS to AR signaling and castration resistance needs to be fully elucidated. Finally, single-cell analysis in the human prostate under ADT/ARSI conditions will provide additional insight into the findings presented here.

## ACKNOWLEDGEMENTS

Work in the authors’ lab was supported, in part, by grants from the U.S National Institutes of Health (NIH) R01CA237027, R01CA240290, R21CA237939, and R21CA218635, grants from the U.S Department of Defense (DOD) PC220137 and PC220273, the PCF (Prostate Cancer Foundation) Challenge Award, and by RPCCC and NCI center grant P30CA016056 (all to D.G.T). This work was further supported in part by the NIH grant R01CA238005 (M.M.S). We acknowledge the assistance from X. Williams and the Atlas Studios in generating the Graphical Abstract and Figure 7F and the Roswell IT group, including K. Quinn, P. Whalen and J. Lee for helping to establish the public facing web portal for our datasets. We thank all other Tang lab members for helpful discussions throughout this project. We apologize to colleagues whose work cannot be cited due to space constraint.

## AUTHOR CONTRIBUTIONS

J.S.K and D.G.T. conceived the project and designed all studies in the project. J.S.K., with the assistance from A.T., R.K., P.K.S., and J.L., performed most experiments. J.W., M.L., S.P., H.Y., S.L., and T.L. are professional bioinformaticians and conducted data acquisition, analysis and interpretation associated with the scRNAseq and scATAC experiments. Y.J. and X.L. conducted some further bioinformatics analysis linking mouse data to human PCa. H.Y. was involved in experimental design of some *in vivo* studies and in all statistical analysis. Q.C and J.H. provided their original scRNAseq data in human PCa samples and participated in our re-mapping and data analysis. J.L.W. and M.M.S. provided the *Clu*;Tomato and *Nkx*;EYFP mouse lines, respectively, and were involved in some experimental design and data analysis. I.P. and G.C. were involved in the initial project conception and offered critical clinical insight. J.S.K. wrote the manuscript draft, and J.S.K and D.G.T finalized the manuscript.

## STAR Methods

### RESOURCE AVAILABILITY

#### Lead Contact

Further information and requests for resources and reagents should be directed to and will be fulfilled by the lead contact, Dean G. Tang.

#### Materials availability

No new mouse models or reagents were generated for these studies. Models used are available through MTAs associated with the original authors and their respective institutes. Where indicated, mice can be purchased from Jackson Laboratory. All reagents are commercially available.

#### Data and code availability

- Single-cell RNA-seq and ATAC-seq data have been deposited at Gene Expression Omnibus (GEO) and are publicly available as of the date of publication. Accession numbers are listed in the key resources table. This paper also analyzes existing, publicly available data. Accession numbers for public datasets are listed in the key resources table. Microscopy data reported in this paper will be shared by the lead contact upon request.
- All original code is publicly available as of the date of publication. Digital Object Identifiers (DOIs) are listed in the key resources table.
- Any additional information required to reanalyze the data reported in this paper is available from the lead contact upon request.

### EXPERIMENTAL MODEL AND SUBJECT DETAILS

#### Mouse Models

All animal related experimental procedures were approved by Roswell Park Comprehensive Cancer Center (RPCCC) Institutional Animal Care and Use Committee (IACUC). All mice used in these studies were male and between the ages of 7 and 13 weeks of age, as indicated in experimental outlines. Only males develop prostate glands, and hence female mice were excluded from these studies. All mice were housed under standard conditions in groups of 2 to 4 with littermates, fed a standard mouse diet, and provided drinking water *ad libitum*.

C57BL/6J mice were purchased from Jackson laboratory (RRID:IMSR_JAX:000664) and used for both single-cell sequencing studies and the testing of therapeutic interventions during prostate castration/regeneration. *Clu*-CreER(T2) mice^37^ were kindly provided by Dr. Jeffery Wrana. *Clu*- CreER(T2) mice were crossed with B6.Cg-*Gt(ROSA)26Sor^tm^*^14^(CAG–tdTomato)*^Hze^*/J mice (RRID:IMSR_JAX:007914) to generate the *Clu*;tdTomato model. The *Nkx3.1*-CreER(T2) model^38^ (B6;129S-*Nkx3-1^tm^*^3^(cre)*^Mms^*/Nci) was kindly provided by Dr. Michael Shen^38^, but can also be acquired from the NCI Mouse Repository (#01XBP). *Nkx3.1*-CreER(T2) mice were crossed with B6.129X1- *Gt(ROSA)26Sor^tm^*^1^(EYFP)*^Cos^*/J mice (RRID:IMSR_JAX:006148) to generate the *Nkx3.1*;EYFP model. Finally, B6N.129S6(Cg)-*Krt5^tm^*^1^.^1^(cre/ERT2)*^Blh^*/J (RRID:IMSR _JAX :029155) were crossed with tdTomato to create the *Krt5*;tdTomato model. All model colonies were maintained as heterozygous for the Cre driver, but homozygous for tdTomato/EYFP, with Cre always coming from the male mouse.

### METHOD DETAILS

#### Prostate castration and regeneration assays

Male wild type C57BL/6 mice (RRID:IMSR_JAX:000664) were purchased from Jackson laboratory at 6 weeks of age. At 8 weeks of age, mice were either left intact or castrated via approved IACUC methods. After 4 weeks, a cohort of castrated mice was implanted subcutaneously with silastic tubing containing 14 mg of testosterone (T) to induce prostate regeneration. Intact and castrated mice were euthanized at 12 weeks of age. Mice undergoing T induced prostate regeneration were euthanized 96 hours post testosterone implantation. At the time of euthanasia, the genitourinary (GU) tract was dissected out for prostate dissection and downstream analysis.

To determine whether prostate regeneration could be affected by therapeutic intervention, mice underwent castration/regeneration as described above. However, mice (N=8/treatment group) were also started on treatment 2 days post-castration with either vehicle [10% vol/vol DMSO (#D8418, MilliporeSigma), 40% vol/vol PEG300 (#202371, MilliporeSigma), 5% vol/vol Tween80 (#P4780, MilliporeSigma), mixed in distilled water] or drugs of interest, including Panobinostat (10mg/kg QOD, 3x/week; IP injection; #HY10224, MCE), XAV939 (2.5mg/kg QID, 1x/every 4 days; IP injection; #HY- 15147, MCE), ML-355 (30mg/kg QOD, 3x/week; IP injection; #HY-12341, MCE), Bortezomib (1mg/kg QD, 1x/week; IV injection; #HY-10227, MCE), Tanzisertib (10mg/kg QD, 5x/week; PO injection; #HY- 15495, MCE), and Capivasertib (100mg/kg QD, 5x/week; PO injection; #HY-15431, MCE)(Figure 6I, Key resource table). DMSO was max 10% but concentration was dependent on drug solubility.

#### Tissue processing and dissociation

Mouse prostates from various conditions were dissected from the GU tract and processed for histological purposes or for tissue dissociation to prepare single cells. For histological purposes prostate tissues were either fixed with 10% neutral buffered formalin (NBF)(#NBF-4-G, AzerScientific), dehydrated with 70% ethanol (#111000200, Pharmco) and paraffin embedded (FFPE) according to standard protocols, or were fixed in fresh 4% paraformaldehyde (PFA)(#15714, EMS) diluted in PBS at 4°C for 4 hours, serially passaged through 10, 20, and 30% sucrose (#J64270.A1, ThermoScientific) for 24 hours each, and subsequently embedded in OCT medium (#3801480, Surgipath). FFPE and OCT tissue blocks were cut at 5 μm and 7.5 μm, respectively, and placed on positively charged glass slides for downstream use and analysis.

Single-cell tissue dissociations were performed as previously described^85^. Briefly, in dissecting medium (DMEM (#10-027-CV, Corning) with 10% (vol/vol) FBS (#10437028, ThermoFisher), 1X glutaMAX (#35050061, ThermoFisher), and 1X penicillin-streptomycin (#15070063, ThermoFisher)) prostates were dissected away from the GU tract into individual lobes from 4-6 independent mice (N=4- 6). Dorsal, lateral, and ventral prostatic lobes were pooled together, and anterior lobes were processed separately. Pooled prostatic tissue was minced and transferred to digestion solution containing 1 mg/ml collagenase (#17100017, ThermoFisher Scientific). Tissue was digested at 37°C for 2 hours with constant inversion. After digestion, tissue was spun down, resuspended in Trypsin-EDTA (#25300120, ThermoFisher), and digested for an additional 5 minutes at 37°C. Trypsin digestion was stopped by addition of dissecting medium supplemented with 500 U DNase I (#10104159001, Millipore-Sigma). Digested tissue was next passed through 18-G and then 20-G syringes to facilitate dissociation. Dissociated tissue was finally passed through a mesh filter with a 40 μm pore size to isolate single cells. All solutions used for tissue digestion/dissociation contained 10 μM Y-27632 dihydrochloride (#1254, Tocris) to prevent anoikis. Cell viability was greater than 80% for all downstream assays.

#### Single-cell RNA sequencing and data processing

##### Single-cell RNA library generation and sequencing

Single-cell RNAseq for dissociated mouse tissues was performed using 10X genomics Chromium Single-Cell 3’ Library & Gel bead Kit V2 (ADLVP study) or the 10X genomics Chromium Single-Cell 3’ Library & Gel bead Kit V3 (AP2 study) according to the manufacturer’s protocol. Briefly, approximately 17,000 viable cells were loaded into each channel of the Chromium Next GEM Chip G to capture 10,000 Gel Beads-in-emulsion (GEMs) per channel. After GEM capture, gel beads are dissolved and cells are lysed into the cDNA master mix containing reverse transcription (RT) reagents. Post RT incubation, GEMs were broken, and first-strand barcoded-cDNA was purified with magnetic beads, followed by 12- cycles of PCR amplification. PCR-amplified barcoded-cDNA was then fragmented and purified with SPRI beads to obtain an average fragment size of 600 bp. Next, cDNA libraries were ligated to sequencing adapters followed by indexing PCR. Resulting libraries were sequenced on the Illumina NovaSeq 6000. Cell-ranger v2.1.0 (10X genomics) was used to demultiplex all mouse runs to FASTQ files, align reads to the mm10 mouse transcriptome and extract cell and UMI barcodes. The pipeline output is a cell-by-gene count matrix, which records the number of UMIs for each gene associated with a particular cell barcode.

##### Data processing

The output from the 10X Genomics Cellranger pipeline (version 2.1.1.1) was used as input for Seurat (version 3.1.1) in R statistical software (version 4.1.1). Low-quality cells, characterized by either low or high unique feature counts (<500) or high mitochondrial counts (>5%), were filtered out from the analysis. The data was then normalized, scaled, and explored using Seurat’s recommended workflow and parameters. Principal component analysis (PCA) was done for dimension reduction and Graph-based clustering was done to identify clusters. Finally, UMAP (Uniform Manifold Approximation and Projection) analysis was performed to visualize cell clusters. Normalized gene expression was then obtained by implementing the “LogNormalize” function in Seurat, using a scaling factor of 10,000, whereby feature counts for each cell are divided by the total number of counts for that cell and then multiplied by the scale factor (10,000). The value is then natural-log transformed using ‘log1p’. Differential expression between corresponding clusters was assessed using the FindMarkers function of Seurat, which uses a Wilcoxon-Rank Sum test (log2 foldchange cutoff 0.25) to identify the unique gene markers for each cluster.

Two approaches were adopted to exclude doublets. First, we used Scrublet to identify and exclude potential doublets^86^. Scrublet assumes a prior probability of 5% for co-encapsulation doublets given our experimental parameters. Second, cells that shared a high-level overlap between two or more lineages (epithelial, stromal, immune) were also excluded from analysis.

#### RNAscope in situ hybridization

Freshly cut FFPE and OCT tissue sections were stained using RNAscope® Multiplex Fluorescent Reagent Kit v2 (ACD, #323110). Staining was performed as recommended by manufacturer (323100- USM/Rev Date:02272019). Briefly, slides were pre-processed based on collection method according to manufacturer’s recommendations, blocked for endogenous peroxidase activity, underwent antigen retrieval, and protease digestion. Indicated RNAscope® probes (see Key Resource Table) were hybridization to tissue sections at 40 °C for 2 hours. Probe signals were amplified by serial incubations with AMP 1, AMP 2, and AMP 3 reagents and developed using HRP mediated signal conjugation of Fluorescein (#NEL741001KT, AkoyaBio), Cy3 (#NEL744001KT, AkoyaBio), and Opal 690 (#FP1497001KT, AkoyaBio) were used for C1, C2, and C3 channels respectively. After staining the sections were mounted with coverslips using Prolong^TM^ Gold Antifade Mountant with DAPI (#P36931, ThermoFisher). Imaging was performed using the Keyence BZ-X710 all-in-one fluorescent microscope.

#### Gene signature enrichment using AUCell

AUCell allows researchers to identify active gene regulatory networks within individual cells^32^. Here, prostate specific gene regulatory networks from various sources (Table S7) were used as inputs into the AUCell function from SCENIC. Briefly, AUCell uses ranked gene expression in a particular cell to calculate the enrichment of a particular gene set as an area under the recovery curve (AUC). Using the ranked gene expression for a cell, AUCell determines whether the input gene set is enriched in the highest expressed genes for any given cell. Therefore, within each cell the AUC represents the proportion of expressed genes for any signature and their relative expression compared to other genes. The AUC scores for a particular signature within each cell were used to generate a binary matrix using algorithm generated cutoffs and then plotted as an overlay on the UMAP projection for each cell.

#### Reference atlas mapping

Reference-based mapping of previously published mouse prostate scRNA-seq datasets^3,8^ was performed against our newly derived mouse prostate single-cell reference. Raw counts for query datasets were obtained from Gene Expression Omnibus (GSE150692, GSE146811) and re-processed using nearly identical Seurat-based workflow as applied to our dataset (described above). Cell annotations were obtained from GEO and shown as reported in the initial publications. Reference-based mapping and label transfer of previously annotated cells were performed using the FindTransferAnchors and MapQuery functions implemented within Seurat v4^34^.

#### Single-cell ATAC sequencing and data processing

##### Single-cell ATAC library generation and sequencing

Single cell ATAC-Seq libraries are generated using the 10X Genomics platform. Isolated nuclei suspensions are treated with Transposase to preferentially fragment open DNA and add partial adapter sequences to the ends of the DNA fragments. Nuclei are loaded into the Chromium Controller (10X Genomics) where they are partitioned into nanoliter-scale Gel Beads-in-emulsion with a single barcode per cell. Libraries are completed by PCR amplification to incorporate remaining Illumina adapter sequences and unique sample indexes. The resulting libraries are evaluated on D1000 screentape using a TapeStation 4200 (Agilent Technologies) and quantitated using Kapa Biosystems qPCR quantitation kit for Illumina. Libraries are then pooled, denatured, and diluted to 300pM with 1% PhiX control library added. The resulting pool is then loaded into the appropriate NovaSeq Reagent cartridge and sequenced on a NovaSeq6000 per the manufacturer’s protocol (Illumina Inc.). Cellranger (10X Genomics) was used to demultiplex all mouse runs to FASTQ files, align reads to the mm10 mouse genome and extract cell and UMI barcodes. The pipeline output is a cell by DNA fragment count matrix, which records the number of UMIs for each DNA fragment associated with a particular cell barcode.

##### Data processing

The output fragment files from the 10X Genomics Cellranger-arc pipeline (version 1.2.0) were used as input for ArchR (version 1.0.1) in R statistical software (version 4.1.1). ArchR^35^ generated a tiled map of read coverage over the whole genome with bin size of 500bps. Low quality cells, characterized by either low counts (<500) or low reads enrichment at transcription start sites (<4%), were filtered out from the analysis. We applied the functions ‘addDoubletScores’ and ‘filterDoublets’ in ArchR to detect and filter doublets. The data was then normalized, scaled, and explored using ArchR’s recommended workflow and parameters. The ArchR IterativeLSI method was used to reduce dimensionality to 30, then ArchR called Seurat graph clustering method to detect clusters with resolution of 0.3. The UMAP (Uniform Manifold Approximation and Projection) analysis were performed to visualize cell clusters. GeneScore, the prediction of gene expression based on the accessibility of regulatory elements in the vicinity of the gene, was added for each gene using ArchR model discussed in ArchR paper^35^. Then imputation through the MAGIC algorithm was used to smooth the gene score matrix and fill in the missing data. In order to call the differentially accessible regions, we first followed ArchR best practice to call the pseudo-bulk replicates, call reproducible peaks for clusters using MACS2^87^, then applied Wilcoxon-Rank Sum test (FDR 0.1, log2 foldchange 0.5) to identify the marker peaks for each cluster. For each cluster, we scanned the motifs in the differentially accessible regions using ArchR’s function addMotifAnnotations based on CisBP database^88^.

##### Integration

We integrated scRNA-seq and scATAC-seq following a 2-step process. First, the unconstrained Integration approach (function addGeneIntegrationMatrix) by ArchR was used to identify general lineage restricted populations. Next, a second iteration of integration was performed constraining basal and luminal lineages identified from unconstrained integration. Using this 2-step process the GeneScore matrix derived from scATAC-seq was integrated with gene expression matrix from scRNA-seq and cell labels were transferred from scRNA-seq to scATAC-seq.

#### Mouse Lineage Tracing Studies

For lineage tracing, the *Clu*;tdTomato, *Nkx3.1*;EYFP, and the *Krt5*;tdTomato models described above were employed. Briefly, pre-castration labeled mice were given tamoxifen by intraperitoneal (IP) injection (2.5mg/mouse; QD for 5 days) at 7 weeks of age. Tamoxifen (#T5648, MilliporeSigma) was prepared by first dissolving in ethanol (#111000200, Pharmco) at 55°C followed by dilution with corn oil (#C8267, MilliporeSigma) to a 25 mg/mL stock solution (5% vol/vol ethanol in corn oil). Mice were castrated at 8 weeks of age, or left intact, and followed for an additional 4 weeks (until 12 weeks of age). At 12 weeks of age, a subset of castrated mice were implanted subcutaneously with silastic tubing containing 14 mg of testosterone to induce prostate regeneration. After 96 hours of regeneration, all mouse cohorts were humanely euthanized, GU tracts were harvested, and prostates were dissected out for processing and downstream analysis. Lineage traced prostates were generally processed by OCT embedding for downstream analysis. Post-castration lineage traced mice were handled in the same manner with one exception, mice were given tamoxifen by IP injection at 11 weeks of age, post-castration and prior to regeneration.

#### Immunostaining, and image analysis

For immunofluorescent staining PFA fixed and OCT embedded tissue sections were used. Briefly, slides were hydrated in PBS and then permeabilized with 0.3% (vol/vol) Triton X-100 (#T9284, MilliporeSigma) in phosphate buffered saline (PBS) for 10 min. After permeabilization slides were washed in PBS and then blocked with 2% (wt/vol) bovine serum albumin (BSA; #130-091-376, MiltenyiBiotec) in PBS for 1 hour at room temperature. Following blocking slides were incubated with primary antibodies (see Key Resource Table) in antibody incubation buffer (1% (wt/vol) BSA, 0.3% (vol/vol) Triton X-100 in PBS) at optimized dilution overnight at 4 °C in a humidified chamber. After primary antibody incubation slides were washed 2X in PBS and then incubated with appropriately conjugated secondary antibody (see Key Resource Table) for 2 hours at room temperature in a humidified chamber. Lastly, slides were washed 2X in PBS and coverslips were mounted with Prolong^TM^ Gold Antifade Mountant with DAPI (#P36931, ThermoFisher).

Whole mount anterior prostate (AP) images were captured using the Keyence BZ-X710 All-in-one fluorescent microscope. Using the advanced observation module 40x images were stitched together to form seamless high-resolution images of the whole AP. For quantitative analysis of lineage tracing models, representative images (N=3) of stained proximal and distal APs from independent lineage traced mice (N=2) were captured using a Leica TCS SP8 confocal microscope. Representative images were opened in ImageJ software to manually count KRT8 and KRT5 positive lineage traced cells per region of interest (ROI). The number of lineage-traced KRT8 and KRT5 positive cells in the distal and proximal glands were plotted as a percentage of the whole using Graph Pad Prism for individual models.

#### CytoTRACE and pseudotime analysis

CytoTRACE is a publicly available resource designed to predict ordered differentiation states from scRNA-seq data^42^. Furthermore, without any prior knowledge CytoTRACE can predict the direction of differentiation. Briefly, using the number of detectably expressed genes per cell, gene count signatures (GCSs) were calculated. GCSs are defined as the geometric mean of the top 200 genes most correlated with gene counts. Next nearest neighbor graphs were created by conversion of the normalized expression matrix into a Markov process capturing the local similarity of cells. Non-negative least squares regression (NNLS) was applied to GCS by using the Markov matrix to represent GCS as a function of transcriptional neighborhoods in the Markov matrix. GCS is iteratively adjusted based on the probability structure of Markov process, and resulting values for each cell are ranked and scaled between 0 and 1 (0, more differentiation; 1, less differentiated). See cytotrace.stanford.edu for details.

Using the predicted differentiation status of luminal cells, a pseudotime trajectory line was drawn using the trajectory function “addTrajectory” in ArchR between least differentiated and most differentiated populations specified by CytoTRACE. We then extracted changes of integrated gene expression from scRNAseq and motif deregulation over our pseudotime trajectory line, with L5 or L1 populations as the root, or time ‘0’ and the most differentiation population as time ‘100’.

#### Ingenuity Pathway Analysis and GSEA

As discussed, differential expression between corresponding clusters was assessed using the FindMarkers function of Seurat. Alternatively, DEGs were generated using the polygonal selection tool and the locally distinguishing gene function in Loupe Browser v6.1.0 for populations of interest. DEG lists were uploaded into ingenuity pathway analysis (IPA)^43^ to determine potential pathway deregulation and upstream regulators of the specified DEG response (Tables S6 and S7). All pathway and upstream regulator analysis was performed according to IPA recommendations and cutoffs. Select DEG lists were further analyzed by gene set enrichment analysis (GSEA) version 4.2.1 using the ‘hallmark’ gene sets in Molecular Signature Database (MsigDB) version 7.5 provided by the Broad Institute^89, 90^ according to user guidelines. The FDR for GSEA is the estimated probability that a gene set with a given normalized enrichment score (NES) represents a false-positive finding.

#### Reprocessing of publicly available datasets

Deposited BAM files from Cheng et al.^48^ were obtained from the sequence read archive (SRA)(#PRJNA699369) and reprocessed using standard pipelines that apply 10x’s ‘bamtofastq’ and ‘cellranger’ tools. Using Loupe Browser v6.1.0, single cells with fewer than 512 and greater than 65,536 UMIs per barcode were removed. Furthermore, single cells with less than 1,024 and greater than 8,192 genes per barcode were filtered. Lastly, cells with greater than 7.5% mitochondrial UMIs were filtered. Overall, 24,142 high quality cells were used for downstream analysis. UMAP clusters were determined using a K-means 10 algorithm. Using original sample designations, globally defined gene expression patterns, and AMACR expression, 9 major epithelial cell subtypes were identified. Using the polygonal selection tool, locally distinguished differentially expressed genes (DEGs) were calculated between castrate resistant (CR) and intact luminal cells for use in ingenuity pathway analysis (See Table S7).

### QUANTIFICATION AND STATISTICAL ANALYSIS

Prostate regeneration studies were analyzed by independent sample *t*-tests (two-sided) at a significance level of 0.05. A sample size of N=8 mice/group in order to detect a 50% block of testosterone effect (the interaction effect) by the treatment (80% power). The statistical suite in GraphPad Prism was used to perform two-sided t-tests on individual drug responses in prostate regeneration assays.

