## Supplemental Information for "Integrated single-cell analysis defines the epigenetic basis of castration-resistant prostate luminal cells"

### Summary of Supplementary Materials

#### Supplementary Figures

**Refers to Figure 1**
Figure S1. Mouse prostate single-cell RNA sequencing summary.
Figure S2. Lineage specific marker gene overlays of scRNAseq UMAP.
Figure S3. Single-cell mouse prostate atlas is a resource for other studies.

**Refers to Figure 2**
Figure S4. Experimental design and summary of integrated scRNA sequencing studies.
Figure S5. Quality control summary of integrated scATAC sequencing studies.

**Refers to Figure 3 and 4**
Figure S6. The castration-resistant L1 population is not derived from pre-existent basal cell populations.

**Refers to Figure 7**
Figure S7. L1 significance to human PCa and prostate response to castration.

#### Supplementary Tables

**Refers to whole study**
Table S1. Statistics of the 2 single-cell omics projects (i.e., the ADLVP and AP2) - 229,794 total cells.

**Refers to Figure 1**
Table S2. scRNA-seq quality control metrics and cluster-specific DEGs for ADLVP study.

**Refers to Figure 2**
Table S3. scRNA-seq quality control metrics and cluster-specific DEGs for AP study.
Table S4. scATAC-seq quality control metrics and cluster-specific peak and GeneScores for AP study.
Table S5. scATAC-seq cluster specific normalized motif enrichment for AP study.

**Refers to Figure 6**
Table S6. scRNA-seq DEG and Ingenuity Pathway Analysis (IPA) results summary.

**Refers to Figure 1 and 7**
Table S7. Summary of gene signatures used in AUCell and IPA results summary from human scRNA-seq analysis.

### Supplementary Figure Legend:

#### Figure S1. Refers to Figure 1. Mouse prostate scRNAseq summary and validation.

**(A)** Sequencing samples were collected as depicted in project schematic.

**(B)** H&E staining of mouse prostate before and after castration.

**(C)** Gross images of the mouse genitourinary (GU) organs and the prostate. Upper left: urogenital sinus tract with seminal vesicles (SV), bladder and prostatic lobes (AP, DP, LP and VP refer to anterior, dorsal, lateral and ventral prostate, respectively) still attached to the urethra. Upper right: the prostate with individual lobes attached to the urethra but with bladder and SV removed). Lower right: individual lobes dissected from urethra. Lower left: depictions of proximal (Prox) versus distal regions of epithelial prostatic ducts in relations to the urethra (U).

**(D)** Quality control violin plots from individual scRNA sequencing samples depicting % mitochondrial transcript content per cell, the number of transcripts per cell, and number of genes per cell.

**(E)** Representative images of *in situ* RNA hybridization verifying marker expression profiles in intact and castrated AP from proximal and distal regions.

**(F,H)** Representative images of *in situ* RNA hybridization from mouse prostate whole mounts.

**(G)** Representative images of *in situ* RNA hybridization verifying marker expression profiles in intact anterior (AP) and ventral prostatic (VP) lobes.

**(I)** Bubble plot projection of the unique marker genes of stromal and immune cell specific cell subtypes.

#### Figure S2. Refers to Figure 1. Lineage-specific marker gene overlays of scRNAseq UMAP.

**(A)** UMAP of mouse prostate single-cell clusters annotated for lobe-specific populations, castration-specific populations, and proximal progenitor populations. CR, castration-resistant.

**(B)** Changes in cell population frequencies based on scRNAseq library subtype.

**(C)** Single-cell cluster annotation comparisons from various mouse prostate single-cell studies.

**(D-O)** Marker gene overlays specific to epithelial populations (D), lobe-specific luminal populations (E), castration-resistant (CR) luminal (L1/L3) and proximal progenitors (L5) (F), L1-specific (G), L5-specific (H), genes previously reported to be enriched in CR stem cells (I), basal cells (J), stromal cells (K), immune cells (L), L3/L5 specific (M), smooth muscle (Sm) cells (N), and vascular endothelial (Ve) cells (O).

#### Figure S3. Refers to Figure 1. Single-cell mouse prostate atlas is a resource for other studies.

**(A)** Individual library samples from Karthaus *et al.*, 2020 were mapped to the reference atlas coordinates generated from ADLVP dataset in Figure 1A. Red boundaries indicate emergence of castration-resistant luminal clusters, while green boundaries indicate the presence of pre-existent castration-resistant proximal luminal clusters.

**(B)** Individual library samples from Crowley *et al.*, 2020 mapped to ADLVP reference atlas coordinates from Figure 1A.

**(C)** Changes in epithelial cluster frequencies across samples from Karthaus *et al.*, 2020.

**(D)** Changes in select luminal cell cluster frequencies across samples from Crowley *et al.*, 2020.

**(E)** Predicted cell score matching between cell clusters identified in Figure 1A and those identified by Karthaus *et al.*, 2020.

**(F)** Representative images of *in situ* RNA hybridization verifying marker expression profiles in intact and castrated AP.

**Figure S4. Refers to Figure 2. Experimental design and summary of integrated scRNA sequencing studies.**

**(A)** Schematic summarizing the animal treatment and collection timepoints for integrated single-cell RNA and ATAC sequencing studies.

**(B)** scRNAseq violin plots for % mitochondrial content, number of unique molecular identifiers (UMIs) per cell, and number of transcripts per cell demonstrating good sample quality. Note samples were filtered as described in methods.

**(C)** Heatmap of cell-cluster specific genes identified from scRNAseq with annotation of lineage-specific markers.

**(D)** Bubble plot of unique marker genes from specific luminal, basal, and mesenchymal cell subtypes.

**(E)** UMAP expression overlays for L7/L10 marker genes.

**(F)** UMAP expression overlays for distinct L5-specific marker genes.

**(G)** UMAP expression overlays for distinct L1-specific marker genes.

**(H)** Individual library samples from Karthaus *et al.*, 2020 reference mapped to the AP2 reference atlas coordinates from Figure 2A. Red boundaries indicate emergence of castration-resistant luminal clusters, while green boundaries indicate the presence of pre-existent castration-resistant proximal luminal clusters.

**Figure S5. Refers to Figure 2. Quality control summary of integrated scATACseq studies.**

**(A)** The % of ATACseq fragments versus fragment size for each sample, and the relationship of insertion site and distance from the TSS demonstrating high quality samples.

**(B)** Similar peak distributions are observed for each cell cluster, with most insertion sites occurring at distal or intronic regions.

**(C)** UMAP of 16 distinct cell clusters from the scATACseq study identified prior to integration with scRNAseq.

**(D)** Hierarchical clustering of insertion peaks (red) and GeneScores (yellow) from the scATACseq by cell type demonstrates unique and common chromatin accessibility across 16 unique cell clusters.

**(E-G)** *De novo* motif sequence enrichment for L7 (E), L5 (F), and L9 (G) specific chromatin peaks ranked by significance. EoB, Enrichment over background.

**(H)** Chromatin accessibility plots for *Pbsn* and *Clu* demonstrate cluster-specific regulation of unique genes. Regions A and B represent sites of high co-accessibility between ATAC and RNA seq profiles.

**Figure S6. Refers to Figure 3 and 4. The castration-resistant L1 population is not derived from pre-existent basal cell populations.**

**(A)** Representative images of proximal and distal regions of *pre-castration* labeled *Clu*;Tom APs under intact, castrated, and regenerated conditions.

**(B)** Representative images of proximal and distal regions of *post-castration* labeled *Clu*;Tom APs under castrated and regenerated conditions.

**(C)** Representative images of proximal and distal regions of *pre-castration* labeled *Nkx3.1*;EYFP APs under intact, castrated, and regenerated conditions.

**(D)** Representative images of longitudinal (with proximal prostate at the bottom) whole-mount sections of the AP from *Krt5*-CreER(T2);tdTomato mice labeled *pre-castration*, and collected under either intact, castrated, or regenerated conditions.

**(E)** Representative images of *pre-castration* labeled *Krt5*;Tom APs double stained for KRT8 (AF-488) and KRT5 (AF-647) identifying luminal and basal cells labeled by Krt5;Tom in proximal and distal regions of intact, castrated, and regenerated prostates.

**Figure S7. Refers to Figure 7. L1 significance to human PCa and prostate response to castration.**

**(A)** Changes in candidate gene expression from Rajan dataset pre-/post-ADT.

**(B-C)** Changes in candidate gene expression from Sharma dataset pre-/post-ADT.

**(D)** Scatter plot of IPA predicted TF deregulation in the Rajan dataset compared to predicted TF deregulation in L1. Activation score color and -Log10(p-value) bubble size are indicative of the L1 population.

**(E)** Scatter plot of IPA predicted Drug deregulation in the Rajan dataset compared to predicted Drug deregulation in L1. Activation score color and -Log10(p-value) bubble size are indicative of the L1 population.

**(F)** UMAP of re-analyzed single-cell data from Cheng et al., 2022 overlayed with original sample ID.

**(G)** UMAP expression overlay for *AMACR* in the Cheng et al., 2022 dataset indicating benign versus cancerous prostate single cells.

**(H)** UMAP expression overlays for select markers from the Cheng et al., 2022 dataset.

**(I)** Heatmap of the top DEGs by cluster from Cheng et al., 2022.

**Table S1. Statistics of the 2 single-cell omics projects (i.e., the ADLVP and AP2) - 229,794 total cells.**

| ADLVP[a] Project | Expected | Pre-processing # | Post-processing # |
|---|---|---|---|
| AP[b] Intact | 20,000 | 26,571 | 22,219 |
| AP Castrated | 20,000 | 33,198 | 22,638 |
| DLVP[c] Intact | 40,000 | 65,873 | 45,677 |
| DLVP Castrated | 40,000 | 49,127 | 41,653 |
| AP2 Project | | | |
| AP Intact (RNAseq) | 20,000 | 23,273 | 19,786 |
| AP Castrated (RNAseq) | 20,000 | 18,822 | 16,929 |
| AP Reg[d] (RNAseq) | 20,000 | 17,979 | 14,487 |
| AP Intact (ATACseq) | 20,000 | 40,618 | 14,068 |
| AP Castrated (ATACseq) | 20,000 | 21,025 | 17,794 |
| AP Reg (ATACseq) | 20,000 | 22,358 | 14,543 |
| Sum Total | 240,000 | 318,844 | 229,794 |

[a]Anterior, Dorsal, Lateral, Ventral Prostate (ADLVP).
[b]Anterior Prostate (AP).
[c]Dorsal, Lateral, Ventral Prostate (DLVP).
[d]Regenerated (Reg).

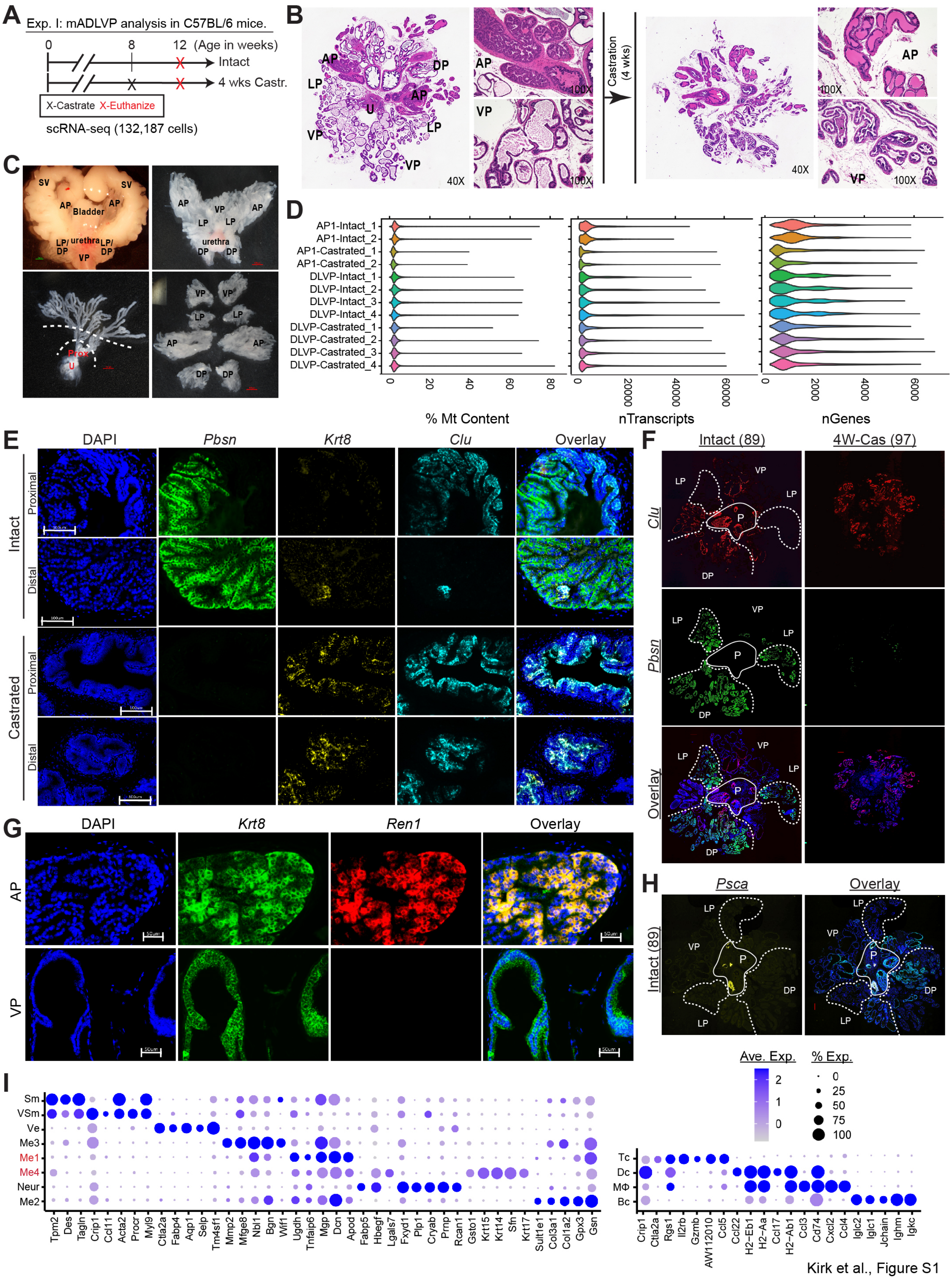

Kirk et al., Figure S1

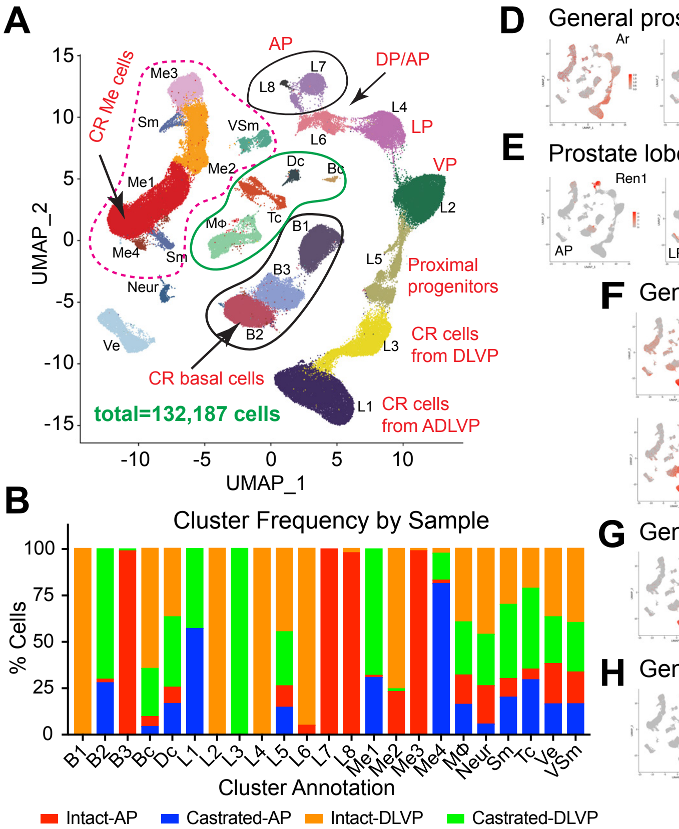

C

**Nomenclature comparison of cell populations**

| Study | Kirk, 2023 | Karthaus, 2020 | Crowley, 2020 | Joseph, 2020 | Guo, 2020 |
|---|---|---|---|---|---|
| Total Cell # | 183, 389 | 88, 395 | 13, 429 | 49, 658 | 45, 503 |
| Cell Type | | | | | |
| Luminal | L1 | | | | |
| | L2 | | LumV | VP Lum | Luminal A |
| | L3 | | | | |
| | L4 | | LumL | DLP Lum | Luminal B |
| | L5 | L2 | LumP | Ur Lum | Luminal C |
| | L6 | | LumD | DLP Lum | Luminal B |
| | L7 | L1 | LumA | AP Lum | Luminal B |
| | L8/L9/L10 | | | | |
| | | L3 | | | |
| | | | PrU | | |
| Basal | B1 | Basal | Basal | Basal | Basal |
| | B2 | | | | |
| | B3 | | Basal | | |
| Mesenchymal | Me1 | | | | |
| | Me2 | M2 | Stromal | Stroma | Stromal |
| | Me3 | M1 | Stromal | Stroma | Stromal |
| | Me4/Me5 | | | | |

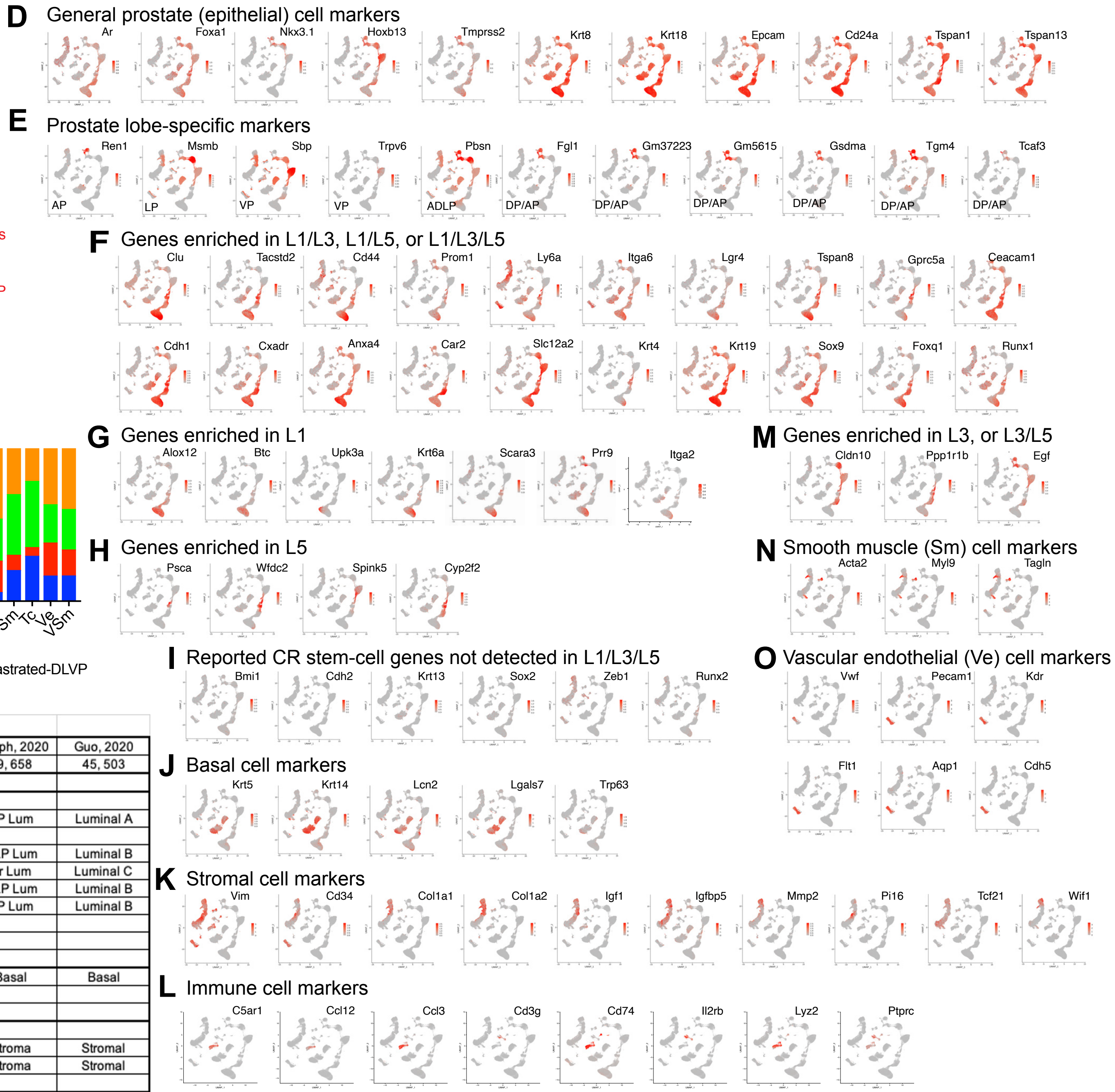

Kirk et al., Figure S2

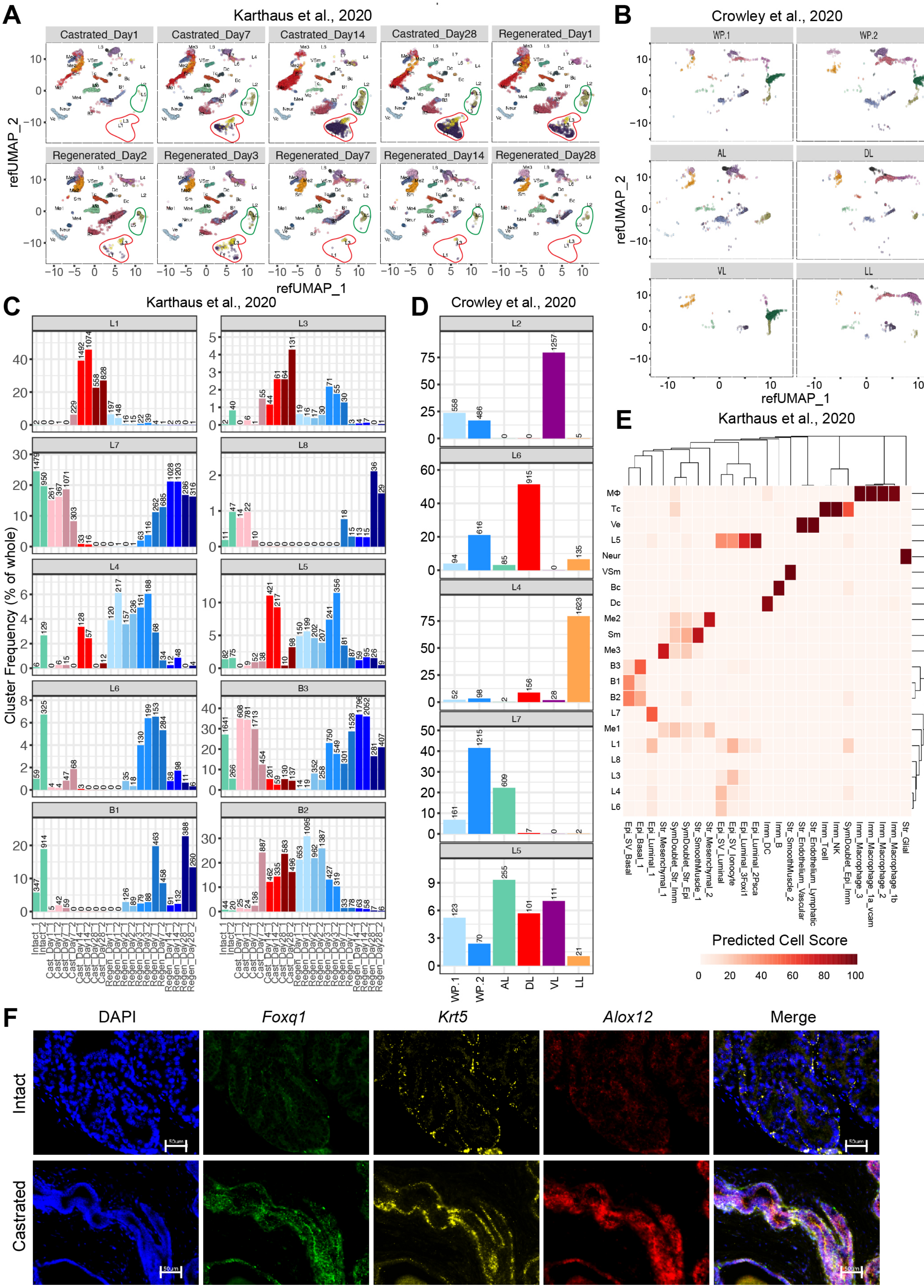

Kirk et al., Figure S3

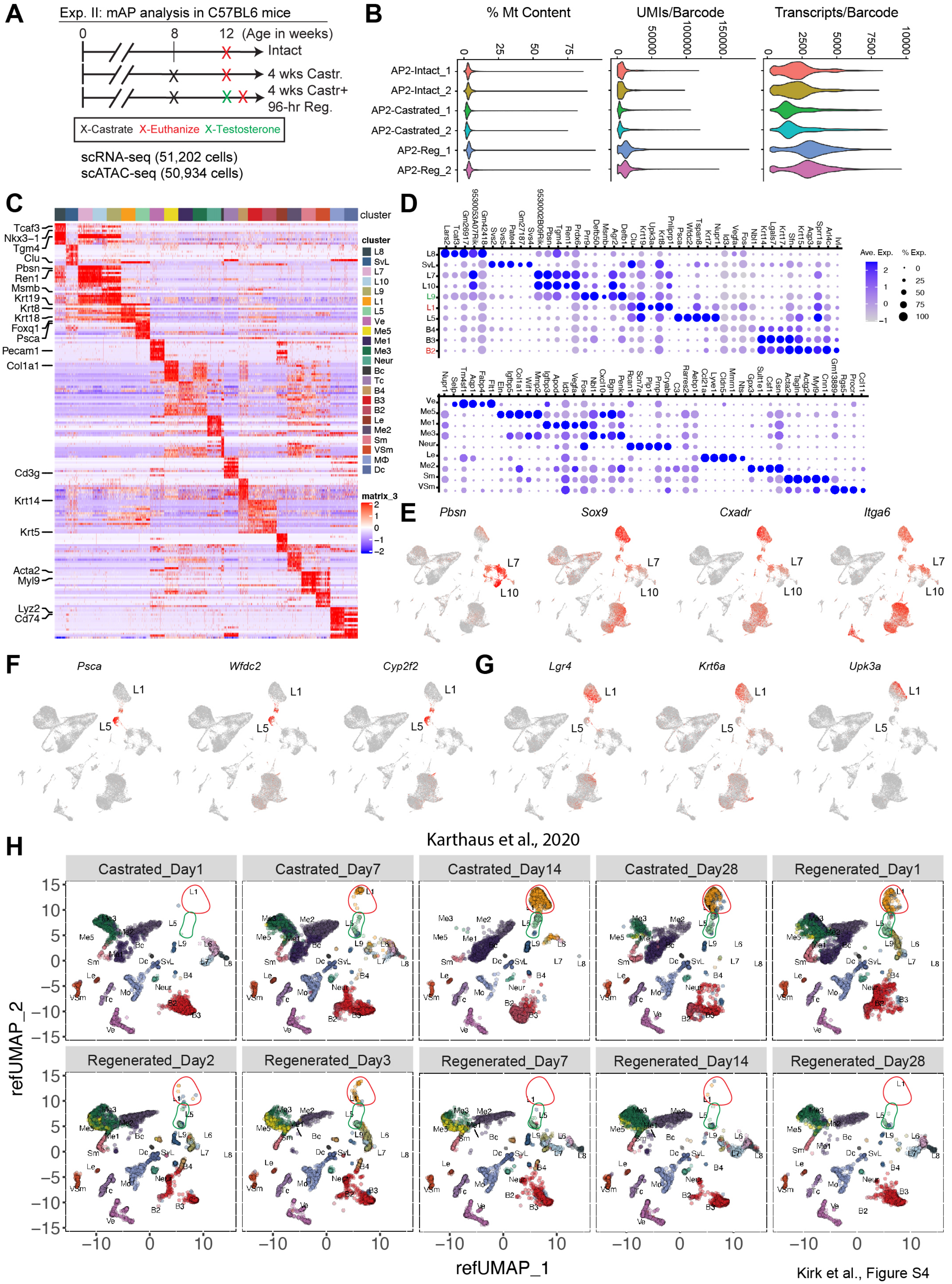

Kirk et al., Figure S4

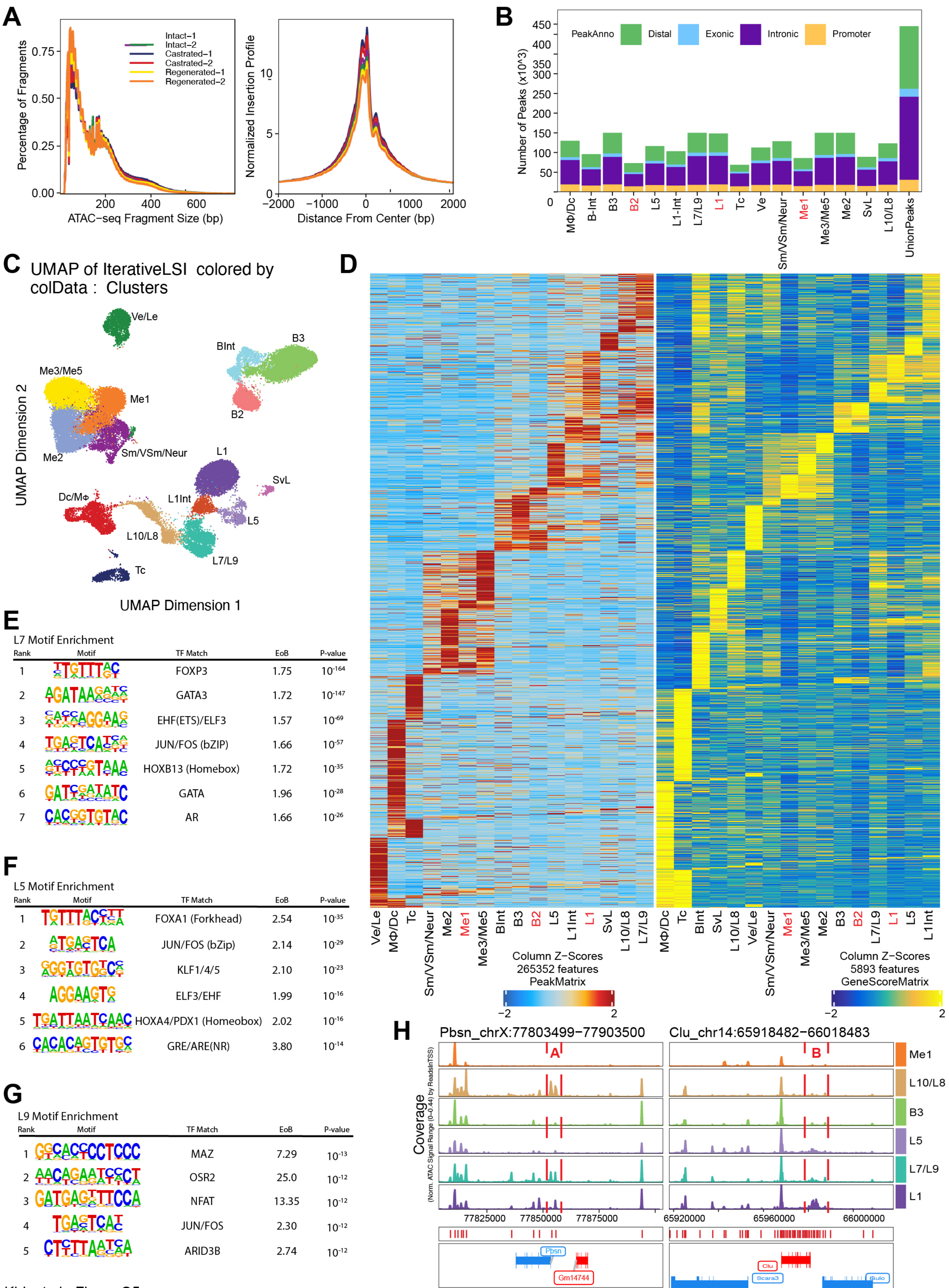

Kirk et al., Figure S5

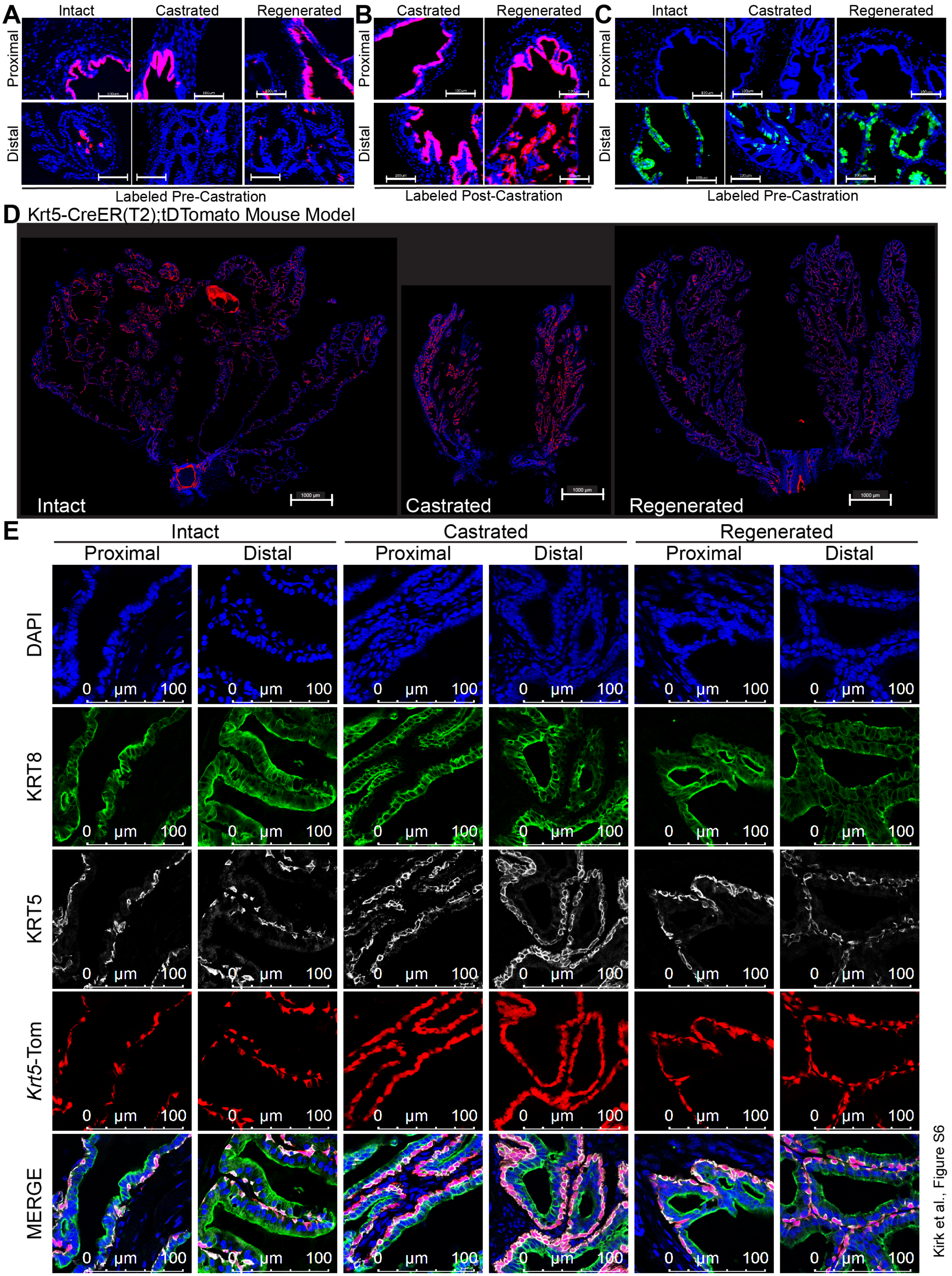

Kirk et al., Figure S6

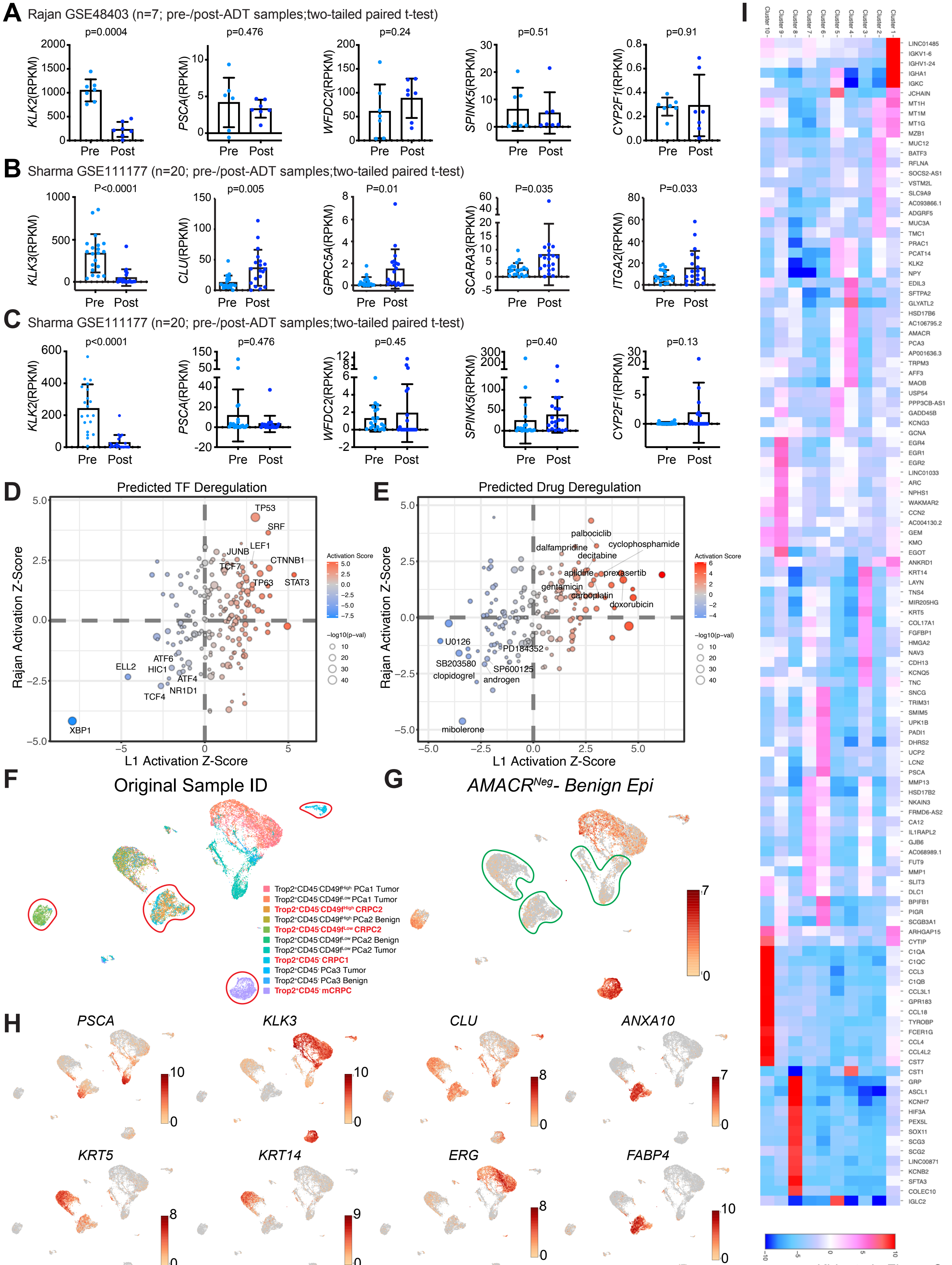

Kirk et al., Figure S7
